# Dissection of a complex disease susceptibility region using a Bayesian stochastic search approach to fine mapping

**DOI:** 10.1101/015164

**Authors:** Chris Wallace, Antony J Cutler, Nikolas Pontikos, Marcin L Pekalski, Oliver S Burren, Jason D Cooper, Arcadio Rubio García, Ricardo C Ferreira, Hui Guo, Neil M Walker, Deborah J Smyth, Stephen S Rich, Suna Onengut-Gumuscu, Stephen J Sawcer, Maria Ban, Sylvia Richardson, John A Todd, Linda S Wicker

**Author notes:** Current address: Department of Chemical Engineering and Biotechnology, University of Cambridge, Cambridge, CB2 3RA, UK. These authors contributed equally (Sylvia Richardson, John A Todd, Linda S Wicker).

## Abstract

Identification of candidate causal variants in regions associated with risk of common diseases is complicated by linkage disequilibrium (LD) and multiple association signals. Nonetheless, accurate maps of these variants are needed, both to fully exploit detailed cell specific chromatin annotation data to highlight disease causal mechanisms and cells, and for design of the functional studies that will ultimately be required to confirm causal mechanisms. We adapted a Bayesian evolutionary stochastic search algorithm to the fine mapping problem, and demonstrated its improved performance over conventional stepwise and regularised regression through simulation studies. We then applied it to fine map the established multiple sclerosis (MS) and type 1 diabetes (T1D) associations in the IL-2RA (CD25) gene region. For T1D, both stepwise and stochastic search approaches identified four T1D association signals, with the major effect tagged by the single nucleotide polymorphism, rs12722496. In contrast, for MS, the stochastic search found two distinct competing models: a single candidate causal variant, tagged by rs2104286 and reported previously using stepwise analysis; and a more complex model with two association signals, one of which was tagged by the major T1D associated rs12722496 and the other by rs56382813. There is low to moderate LD between rs2104286 and both rs12722496 and rs56382813 (*r*^2^ ≃ 0.3) and our two SNP model could not be recovered through a forward stepwise search after conditioning on rs2104286. Both signals in the two variant model for MS affect CD25 expression on distinct subpopulations of CD4+ T cells, which are key cells in the autoimmune process. The results support a shared causal variant for T1D and MS. Our study illustrates the benefit of using a purposely designed model search strategy for fine mapping and the advantage of combining disease and protein expression data.

## Introduction

Genome-wide association studies have been very successful at identifying disease-associated variation by exploiting linkage disequilibrium (LD), meaning that only a subset of markers need be surveyed to detect association. However, this same LD, particularly when combined with the presence of multiple disease causal variants in relative proximity, makes disentangling the specific causal variants difficult. Mapping the likely causal variants in regions associated with complex traits as precisely as possible is becoming increasingly important for two reasons. Firstly, as detailed chromatin annotations become available, more precise maps of probable causal variants will allow researchers to fully exploit these resources through integrative analyses targeted at identifying the specific genes, molecules and cells underlying disease association [1]. Secondly, the functional studies which are ultimately required to confirm causal mechanisms are laborious and need to focus, from the outset, on the smallest yet most complete set of plausible causal variants.

It is widely recognised that the most associated single nucleotide polymorphism (SNP) in a region is not necessarily the causal variant [2], yet attempts at fine mapping disease signals typically proceed by successive conditioning on the most associated SNPs, a form of stepwise regression that may produce incorrect results, particularly when causal variants are correlated, i.e. in linkage disequilibrium (LD) [3]. Bayesian methods have been used for fine mapping association signals in dense and imputed genotyping data and generating credible sets of variants that are likely to contain the causal variant, analogous to credible intervals for odds ratios [4]. However, these approaches typically make the simplifying assumption that exactly one causal variant exists at any individual region [4] because accounting for multiple causal variants leads to exponential increase in the number of models that need to be considered with the number of causal variants considered. Bayesian methods that summarise evidence across SNPs in a region to assess either enrichment of signals in chromatin states [5] or colocalisation between association signals for different traits [6] make a similar single causal variant assumption. As stepwise approaches often obtain evidence for additional, independent association signals in a region [7-10], this assumption is unrealistic. One exception is BimBam [11] which can fit multi SNP models, but considers all possible models up to a specified maximum number of causal SNPs. As the number of potential models grows exponentially, BimBam is limited to regions with relatively few SNPs for computational reasons.

Monte Carlo methods can avoid limitations on the number of causal variants by sampling the model space rather than visiting all possible models. Here we adapt a Bayesian evolutionary stochastic search algorithm, GUESS [12, 13], to the fine mapping problem. This method, and its fast computational implementation, is tailored to efficiently explore the multimodal space created by multiple SNP models. However, the very dense SNP map that is required for fine mapping leads to extreme LD, which presents two specific challenges for GUESS. The first is that SNPs in extremely tight LD can cause numerical instability in model fitting, so we use minimal tagging to explore the model space and then expand all the tag models initially selected by GUESS (Fig. 1).

**Figure 1.**
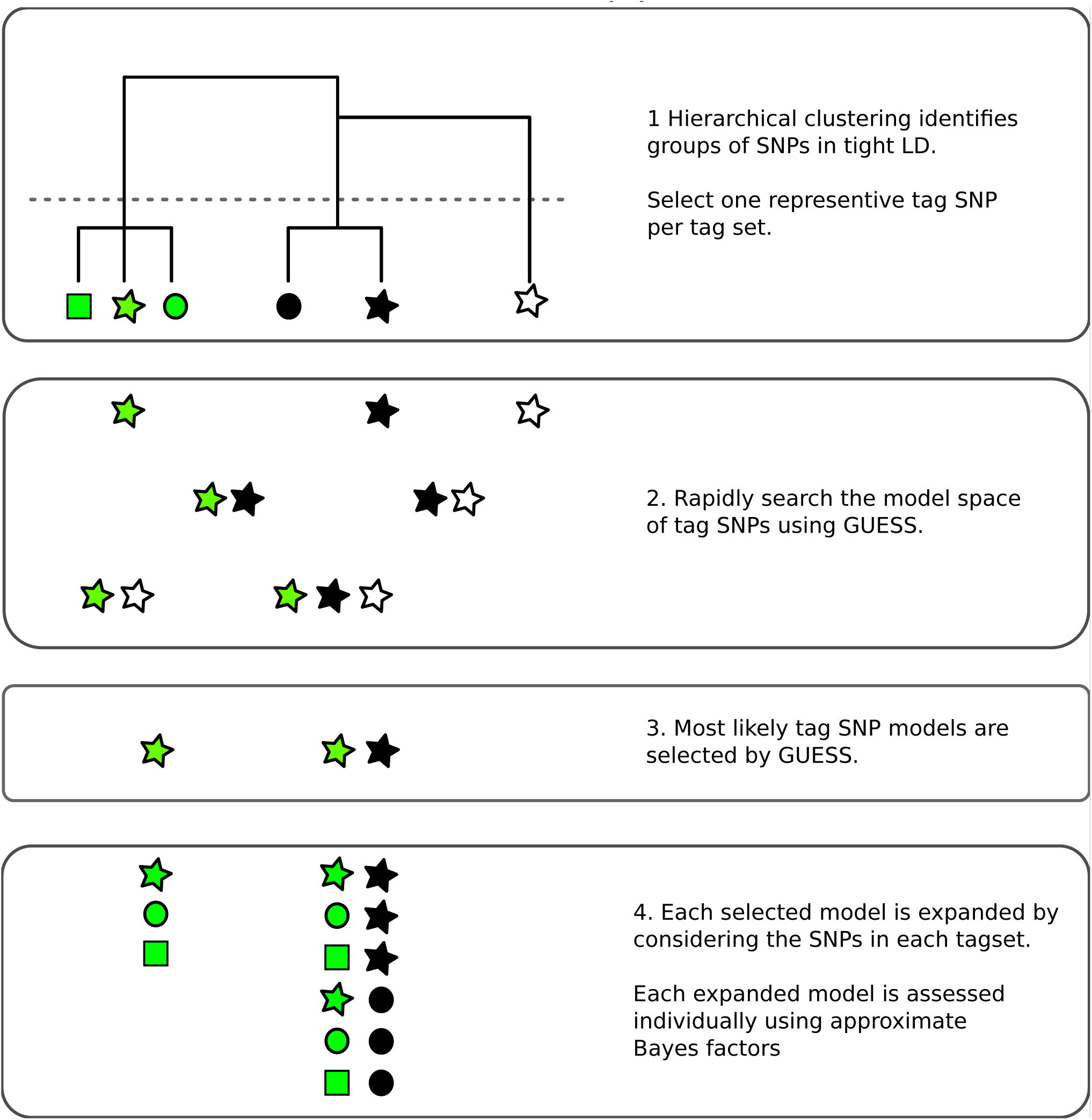
Overview of the fine mapping tailored stochastic search strategy in GUESSFM. 1. SNPs are clustered based on genotype data. Tagging is used to remove cases of extreme LD (*r*^2^ > 0.99) by selecting one SNP from each cluster (“tag set”), that which is in highest average *r*^2^ with all other SNPs. 2. All possible models that can be formed from the tag SNPs may be considered by GUESS. Here, all seven possible models are considered but, in practice, with larger numbers of tags than shown here, GUESS employs a stochastic search strategy to consider only a subset of models, prioritising those with greatest statistical support. 3. GUESS selects the most likely models amongst those it has visited. Here, it selects two of the seven, but in larger data sets we retain the 30,000 most likely. 4. Each of these selected models is expanded by considering all possible substitutions of tags by other members of their tag set. Each expanded model is then assessed again individually, using an approximate Bayes factor [24].

Second, posterior support is diluted across SNPs in tight LD, potentially preventing direct inference on the importance of individual SNPs. We therefore use posterior model probabilities and patterns of LD to define sets of SNPs which have strong joint posterior support for the hypothesis that one member of the set is causal for the trait. These are analogous to the credible sets generated in the Bayesian fine mapping framework which assumes a single causal variant per region [4], but allow for multiple causal variants.

Our adaptions around GUESS are available in an R package, GUESSFM (https://github.com/chr1swallace/GUESSFM). We used GUESSFM to fine map the association of multiple sclerosis (MS) and type 1 diabetes (T1D) to an established susceptibility region for immune-mediated diseases on chromosome 10p15 (Figure S1), which contains the candidate causal gene *IL2RA. IL2RA* encodes a subunit CD25 (IL-2RA) of the interleukin 2 (IL-2) receptor that is essential for the high affinity binding of IL-2 [14]. The region has provided some prime examples of evolving genetic research. Initially associated with T1D susceptibility using a candidate gene and tag SNP approach [15], further studies revealed association to other autoimmune diseases, including MS [16] and rheumatoid arthritis [17], and demonstrated that multiple genetic markers were needed to explain the T1D association [18]. Genotype to phenotype studies have demonstrated that disease-associated variants in the region also associate with *IL2RA* mRNA expression and CD25 protein expression on the surface of naive and memory CD4+ T cells [19-21] and sensitivity of memory CD4+ T and activated T regulatory cells to IL-2 [22].

## Results

The use of different genotyping panels for different diseases has presented a challenge for comparative analysis across diseases. Here, we refined the T1D and MS association signals in the associated region on chromosome 10p15 by taking advantage of a unified dense SNP map provided by the ImmunoChip, a custom Illumina 200K Infinium High-Density array covering 186 distinct autoimmune susceptibility risk regions [7,23]. We used genotypes on a total of 52,637 samples (Table S1), and supplemented the ImmunoChip variants by imputation from the 1000 Genomes Project data. This gave a total of 667 SNPs and small indels with minor allele frequency > 0.005 in controls for analysis after genotype quality control.

To allow for the effects of LD, traditional conditional regression is often supplemented by identifying subsets of SNPs with some minimum LD (e.g. *r*^2^ *>* 0.8) with individual SNPs selected by a forward stepwise procedure, within which it is suggested the causal variant may lie. We compared our proposed approach to traditional conditional regression by simulating phenotypes conditional on multiple “causal variants” selected randomly from within these genetic data, and confirmed that the proposed approach both recovered a higher proportion of the true effects and simultaneously detected fewer false positive results (Figure 2). GUESSFM requires specification of the *a priori* expected number of causal variants in a region, *n*_*exp*_. We considered either setting *n*_*exp*_ = 3 or *n*_*exp*_ = 5, and found the performance decreased when *n*_*exp*_ was less than the true value, but not when it was greater than the true value, suggesting that for recovery of the correct causal variants it is better to set *n*_*exp*_ higher rather than lower. We also considered three regularised regression approaches: the lasso, the elastic net and the group lasso. The ROC curves of those methods (for decreasing penalties) are intermediate to the stepwise and stochastic GUESSFM search approaches (Figure 2). However, at traditional optimization thresholds (minimising the ten fold cross validation), regularized regression approaches were extremely anti-conservative, with a very high false discovery rate, compared to stepwise (stopping at *p <* 10^-8^ or *p <* 10^-6^) or GUESSFM for suitable thresholds on posterior probabilities (Table S2). Picking a regularisation parameter according to the number of predictors included (three or five) produced a more competitive false discovery rate, but discovery rates were still lower than with GUESSFM (Table S2).

**Figure 2.**
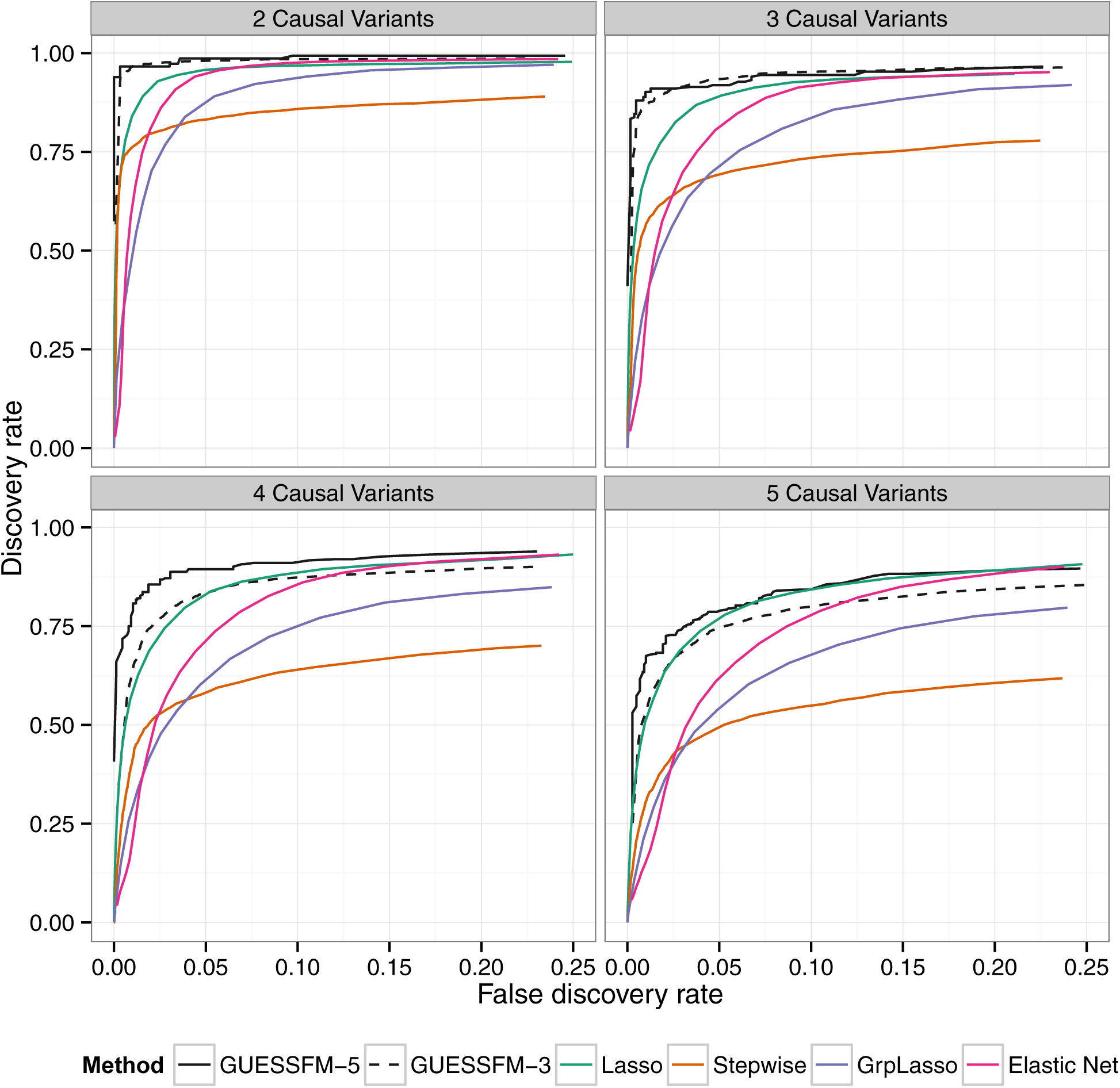
Comparison of of several multivariate methods for fine mapping using simulated data. We simulated quantitative phenotype data with between two and five causal variants using genotype data from the T1D dataset for the *IL2RA* region. The simulated data sets were analysed using forward stepwise regression, GUESSFM, the lasso, the group lasso and the elastic net. GUESSFM produces credible sets for each variant chosen using the snp.picker algorithm described in Materials and Methods. We defined pseudo “credible sets” for the other approaches as the set of SNPs with *r*^2^ > 0.8 with a selected SNP. We calculated the discovery rate (the proportion of causal variants within at least one credible set, y axis) and false discovery rate (proportion of detected variants whose credible sets did not contain any causal variant, × axis) at different thresholds for the stepwise *p* value, the group marginal posterior probability of inclusion (gMPPI) for GUESSFM and the regularization parameter(s) across simulated datasets (see Methods for details). GUESSFM-3 and GUESSFM-5 refer to GUESSFM run with a prior expectation of three or five causal variants per region, respectively. Results are averaged over 1000 replicates.

We applied our approach to the MS and T1D data sets. Posterior support for the number of SNPs required to model the association is strongly peaked, favouring a two-SNP model in MS and a four SNP model in T1D (Figure S2). We summarised inference for SNPs over multiple models by considering the support for a SNP group according to the sum of posterior probabilities (PP) over all models containing a SNP from that group. We term this the group marginal posterior probability of inclusion (gMPPI), which can be understood as the probability that one SNP in the group is causal for the trait of interest. For T1D, there is high confidence (gMPPI > 0.8) for four SNP groups that we denote according to their order in Figure S3: A (indexed by rs12722496), C (rs11594656), E (rs6602437) and F (rs41295159). As the T1D data have previously been analysed using stepwise regression [23], we compared our results to those found by forward stepwise analysis of the same data (Table 1). We saw a direct correspondence in terms of LD (*r*^2^ > 0.6) between the SNPs identified by the two approaches (Table 2). However, models found by GUESSFM had larger log likelihoods for a given number of SNPs, indicating that these models offered a better explanation of T1D-associated genetic variation in the region.

**Table 1.** Comparison of results from stochastic GUESSFM and stepwise searches. *p* values are shown for the stepwise search and compare the listed model to the model above (or the null model, for single SNPs). Bayesian Information Criterion (BIC) is shown for stepwise models and index models found through GUESSFM search.

| Model | p | BIC |
| --- | --- | --- |
| *T1D Stepwise* |  |  |
| rs61839660 | $< 2 \times 10^{-16}$ | -140.10 |
| rs61839660+rs11594656 | $7.64 \times 10^{-12}$ | -177.61 |
| rs61839660+rs11594656+rs12220852 | $1.40 \times 10^{-9}$ | -204.35 |
| rs61839660+rs11594656+rs12220852+rs41295159 | $1.39 \times 10^{-7}$ | -218.83 |
| *T1D GUESSFM* |  |  |
| rs12722496+rs11594656+rs6602437 | — | -207.00 |
| rs12722496+rs11594656+rs6602437+rs41295159 | — | -234.95 |
| *MS Stepwise* |  |  |
| rs2104286 | $< 2 \times 10^{-16}$ | -97.25 |
| rs2104286+rs11256593 | 0.00001437106213 | -105.85 |
| *MS GUESSFM* |  |  |
| rs2104286 (“M2”) | — | -97.25 |
| rs12722496+rs56382813 (“M1”) | — | -111.18 |

**Table 2.** LD (*r*^2^) between SNPs in selected models for T1D using stochastic GUESSFM and stepwise searches. Each GUESSFM search signal has a correspondng SNP found by conditional stepwise regression in moderate to strong LD.

| GUESSFM | Stepwise |  |  |  |
| --- | --- | --- | --- | --- |
|  | rs61839660 | rs11594656 | rs12220852 | rs41295159 |
| rs12722496 | **0.88** | 0.02 | 0.02 | 0.00 |
| rs11594656 | 0.02 | **1.00** | 0.26 | 0.02 |
| rs6602437 | 0.05 | 0.25 | **0.62** | 0.01 |
| rs41295159 | 0.00 | 0.02 | 0.00 | **1.00** |

In contrast, for MS, there were no SNP groups with high gMPPI. We instead saw two distinct models, which together accounted for >80% of the posterior support amongst the 514,476 visited models: either model M1, consisting of SNP groups A (indexed by rs12722496) and D (rs56382813), or model M2 consisting only of SNP group B (rs2104286). SNPs from M1 and M2 were rarely selected together (total PP=0.039 across the visited models). rs2104286 was reported as the single associated SNP in the region in the original MS ImmunoChip analysis after forward stepwise regression [7] (Table 1). rs2104286 is in low to moderate LD with both of the M1 index SNPs (*r*^2^ ≃ 0.3, Table 3).

**Table 3.** LD (*r*^2^) between SNPs in selected models for MS using stochastic GUESSFM and stepwise searches. For the preferred M1 model for MS, there is only weak to moderate LD between SNPs chosen by GUESSFM and conditional approaches.

| GUESSFM | Stepwise |  |
| --- | --- | --- |
|  | rs2104286 | rs11256593 |
| M2: rs2104286 | **1.00** | 0.33 |
| M1: rs12722496 | 0.33 | 0.08 |
| M1: rs56382813 | 0.31 | 0.29 |

Haplotype analysis showed that whilst haplotypes carrying the rs2104286:C allele are protective for MS, so is a less common haplotype carrying the rs2104286:T allele and the protective allele at rs56382813 (Table 4). Indeed, the addition of rs2104286 to a model consisting of the haplotypes formed by M1 was non-significant (p=0.196) whilst the addition of the haplotypes formed by the M1 SNPs to a model consisting of only rs2104286 was significant (*p* = 2.64 × 10^-6^). However, the posterior support favours model M1 over M2 only by a factor of 1.7 and this is dependent on our prior expectation for the number of causal variants: had we expected only a single causal variant in the region, we would have favoured M2 by a factor of 1.7, with M1 being favoured only with a prior expectation of 1.8 or more causal variants (Figure S4). Under any reasonable prior, the posterior support for the two models is so close that we cannot choose between them on statistical evidence alone: we must consider them as plausible alternative explanations for MS association in the region. Note that the M1 SNPs were not within the credible set created under the single causal variant assumption, which consists of rs2104286 alone [7].

**Table 4.**
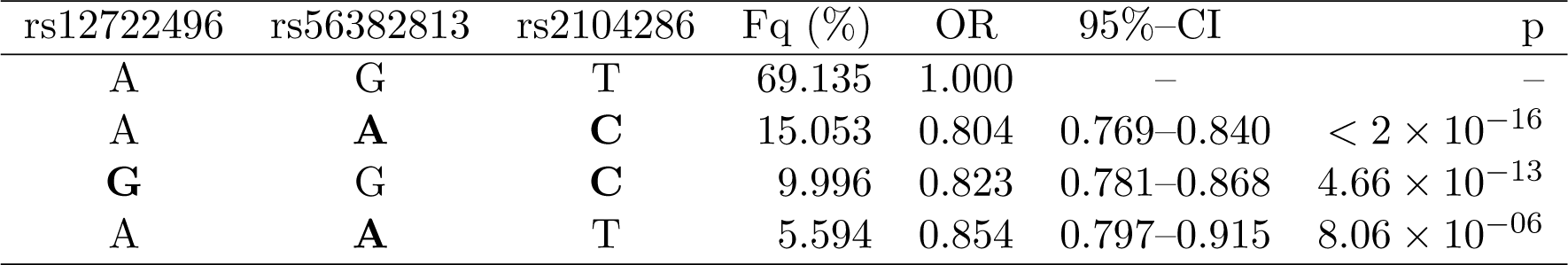
Haplotype analysis of MS association using the index SNPs for the one and two SNP models selected. Minor alleles are indicated by bold font. Fq=haplotype frequency.

We selected all high confidence SNP groups for more detailed exploration (Fig. 3). The T1D signals are located in (1) intron 1 of *IL2RA* - SNP group A, (2) intergenic between *IL2RA* and *RBM17* - C, (3) 5’ of *RBM17* - E, and (4) 5’ of *RBM17* to intron 2 of *PFKFB3* - F. Under the model M1 for MS, SNP group A was also associated with MS, with the same alleles protective for both (Table 5) whilst the second M1 signal (SNP group D) physically overlapped, but was not in LD with, SNPs from group C. Under the model M2, the sole-MS associated SNP (B) is located in intron 1 of *IL2RA,* neighbouring the T1D-associated SNP group A, but there was only weak LD between A and B (*r*^2^ = 0.3).

**Figure 3.**
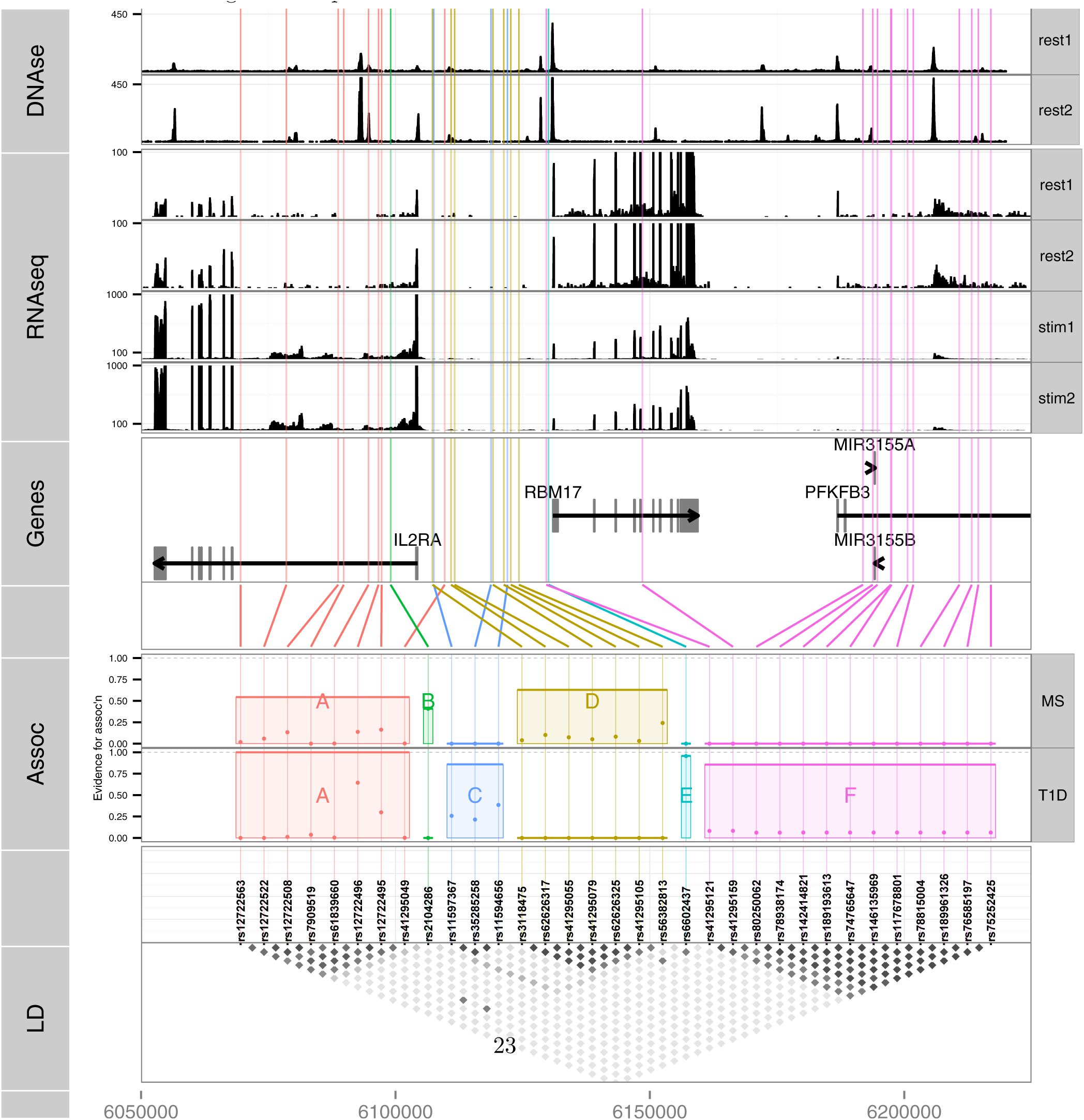
Six sets of SNPs can best explain the association of T1D and MS in the chromosome 10p15 region. **LD**: a heatmap indicating the *r*^2^ between SNPs. Assoc: MPPI for MS and T1D the SNPs in a group, with total MPPI across a SNP group, gMPPI, indicated by the height of the shaded rectangle (see Table 5 for numerical details). SNP groups are labelled by the letters A-F for reference. SNPs in this track are ordered by SNP group for ease of visualisation. Genes: SNPs are mapped back to physical position and shown in relation to genes in the region. RNAseq: read counts in two pooled replicates of resting (“rest1” and “rest2”) and anti-CD3/CD28 stimulated (“stim1” and “stim2”) CD4+ T cells; y axes were truncated to allow visualization of intronic read counts. Note the different limits for resting and stimulated cells, which show greater transcription of all protein coding genes in the region. DNase: DNase hypersensitivity measured in CD4 cells by the Roadmap consortium. Replicate 1 (“rest1”) is R0_01689; replicate 2 (“rest2”) is R0_01736; y axes were truncated again to improve visualization.

**Table 5.** SNPs selected by GUESSFM mapping of MS and T1D associations. SNP groups are labelled A-F for reference. Position is according to hg19; MAF=minor allele frequency in UK controls, OR=odds ratio for minor allele, MPPI=marginal posterior probability of inclusion. SNP groups are divided by horizontal lines, and the group MPPI (gMPPI) is the posterior probability of inclusion of any SNP in the group. We highlight in bold the SNP sets with some support for association with either MS or T1D. The MS model ”M1” consists of groups A and D, the model ”M2” consists of group B.

| Group | SNP | Position | Alleles | MAF | MS |  |  | T1D |  |  |
| --- | --- | --- | --- | --- | --- | --- | --- | --- | --- | --- |
|  |  |  |  |  | Total MPPI | MPPI | OR | Total MMPI | MMPI | OR |
| A | rs12722563 | 6069561 | G>A | 0.107 | **0.544** | **0.019** | **0.858** | **1.000** | **0.000** | **1.000** |
|  | rs12722522 | 6078553 | G>A | 0.098 |  | **0.059** | **0.848** |  | **0.000** | **0.636** |
|  | rs12722508 | 6088743 | A>T | 0.098 |  | **0.134** | **0.845** |  | **0.012** | **0.644** |
|  | rs7909519 | 6089841 | T>G | 0.098 |  | **0.000** | **1.000** |  | **0.037** | **0.641** |
|  | rs61839660 | 6094697 | C>T | 0.089 |  | **0.002** | **0.860** |  | **0.005** | **0.619** |
|  | rs12722496 | 6096667 | A>G | 0.099 |  | **0.139** | **0.845** |  | **0.645** | **0.626** |
|  | rs12722495 | 6097283 | T>C | 0.099 |  | **0.163** | **0.844** |  | **0.300** | **0.628** |
|  | rs41295049 | 6109676 | G>A | 0.087 |  | **0.003** | **0.855** |  | **0.001** | **0.617** |
| B | rs2104286 | 6099045 | T>C | 0.262 | **0.408** | **0.408** | **0.839** | 0.000 | 0.000 | 0.929 |
| C | rs11597367 | 6107534 | A>G | 0.239 | 0.000 | 0.000 | 1.040 | **0.859** | **0.259** | **0.769** |
|  | rs35285258 | 6118770 | C>T | 0.237 |  | 0.000 | 1.039 |  | **0.214** | **0.769** |
|  | rs11594656 | 6122009 | T>A | 0.237 |  | 0.000 | 1.038 |  | **0.386** | **0.767** |
| D | rs3118475 | 6107265 | T>C | 0.191 | **0.628** | **0.040** | **0.811** | 0.000 | 0.000 | 0.941 |
|  | rs62626317 | 6110925 | A>G | 0.191 |  | **0.102** | **0.807** |  | 0.000 | 0.938 |
|  | rs41295055 | 6111622 | C>T | 0.191 |  | **0.075** | **0.808** |  | 0.000 | 0.937 |
|  | rs41295079 | 6119154 | C>T | 0.190 |  | **0.053** | **0.809** |  | 0.000 | 0.942 |
|  | rs62626325 | 6121285 | G>A | 0.176 |  | **0.082** | **0.808** |  | 0.000 | 0.907 |
|  | rs41295105 | 6122674 | T>G | 0.190 |  | **0.033** | **0.812** |  | 0.000 | 0.942 |
|  | rs56382813 | 6124262 | G>A | 0.207 |  | **0.242** | **0.825** |  | 0.000 | 0.907 |
| E | rs6602437 | 6130077 | T>C | 0.433 | 0.000 | 0.000 | 1.018 | **0.956** | **0.956** | **1.187** |
| F | rs41295121 | 6129643 | C>T | 0.009 | 0.000 | 0.000 | 1.000 | **0.857** | **0.082** | **0.541** |
|  | rs41295159 | 6148535 | C>G | 0.009 |  | 0.000 | 0.937 |  | **0.083** | **0.541** |
|  | rs80250062 | 6191884 | T>C | 0.008 |  | 0.000 | 0.941 |  | **0.063** | **0.540** |
|  | rs78938174 | 6193815 | T>C | 0.009 |  | 0.000 | 0.941 |  | **0.063** | **0.540** |
|  | rs142414821 | 6194739 | G>A | 0.008 |  | 0.000 | 0.941 |  | **0.063** | **0.540** |
|  | rs189193613 | 6197361 | G>C | 0.009 |  | 0.000 | 0.941 |  | **0.063** | **0.540** |
|  | rs74765647 | 6197496 | G>A | 0.007 |  | 0.000 | 0.941 |  | **0.063** | **0.540** |
|  | rs146135969 | 6200643 | A>T | 0.008 |  | 0.000 | 0.941 |  | **0.063** | **0.540** |
|  | rs117678801 | 6201774 | C>T | 0.008 |  | 0.000 | 0.941 |  | **0.063** | **0.540** |
|  | rs78815004 | 6210827 | G>A | 0.008 |  | 0.000 | 0.941 |  | **0.063** | **0.540** |
|  | rs189961326 | 6213265 | G>A | 0.008 |  | 0.000 | 0.941 |  | **0.063** | **0.540** |
|  | rs76585197 | 6214560 | G>A | 0.007 |  | 0.000 | 0.941 |  | **0.063** | **0.540** |
|  | rs75252425 | 6217022 | G>A | 0.008 |  | 0.000 | 0.941 |  | **0.063** | **0.540** |

Since we were unable to distinguish between the competing M1 and M2 models for MS using SNP-disease association data alone, we sought corroborating evidence to support SNPs in either model using CD25 flow cytometric expression data. The M2 SNP rs2104286 has been associated with the age-dependent proportion of naïve T cells that express CD25 on their cell surface and rs12722495 (within SNP group A, located in intron 1 of *IL2RA)* with the total expression of CD25 on memory T cells [19,20]. rs12722495 has also been associated with *IL2RA* mRNA expression in T cells, both resting [19] and activated by 48 hour culture with anti-CD3+CD28 [21]. We selected a single tag SNP from each of the credible sets and examined which could best explain CD25 protein expression phenotypes using previously published data from 179 samples [19], plus an additional 30 samples. Both phenotypes were best explained by a single SNP: rs12722495 from group A again showed the strongest association with intensity of CD25 expression on memory T cells (*p* = 5.50 × 10^-10^, Figure S5). For the proportion of naive CD4+ cells that express CD25, the M1 SNP rs41295055 from group D, whose association with CD25 expression was not previously tested, was preferred to the M2 SNP rs2104286 (*p* = 3.45 × 10^-8^ *versus p* = 2.56 × 10^-6^, ABIC=8.43, which is interpreted as “strong” evidence in favour of rs41295055 [24]; Fig. 4), supporting the hypothesis that rs2104286 is not itself functional, but merely tags other functional variants which are causal for MS. The A and D SNP groups coincide with regions in *IL2RA* that contain DNase I sensitivity sites indicating open chromatin available to bind transcription factors under appropriate conditions (Fig. 3). In addition, the existence of RNA-seq reads from resting and stimulated CD4+ T cells within intron 1, where group A SNPs lie (Fig. 3), support the regulatory nature of this region [25].

**Figure 4.**
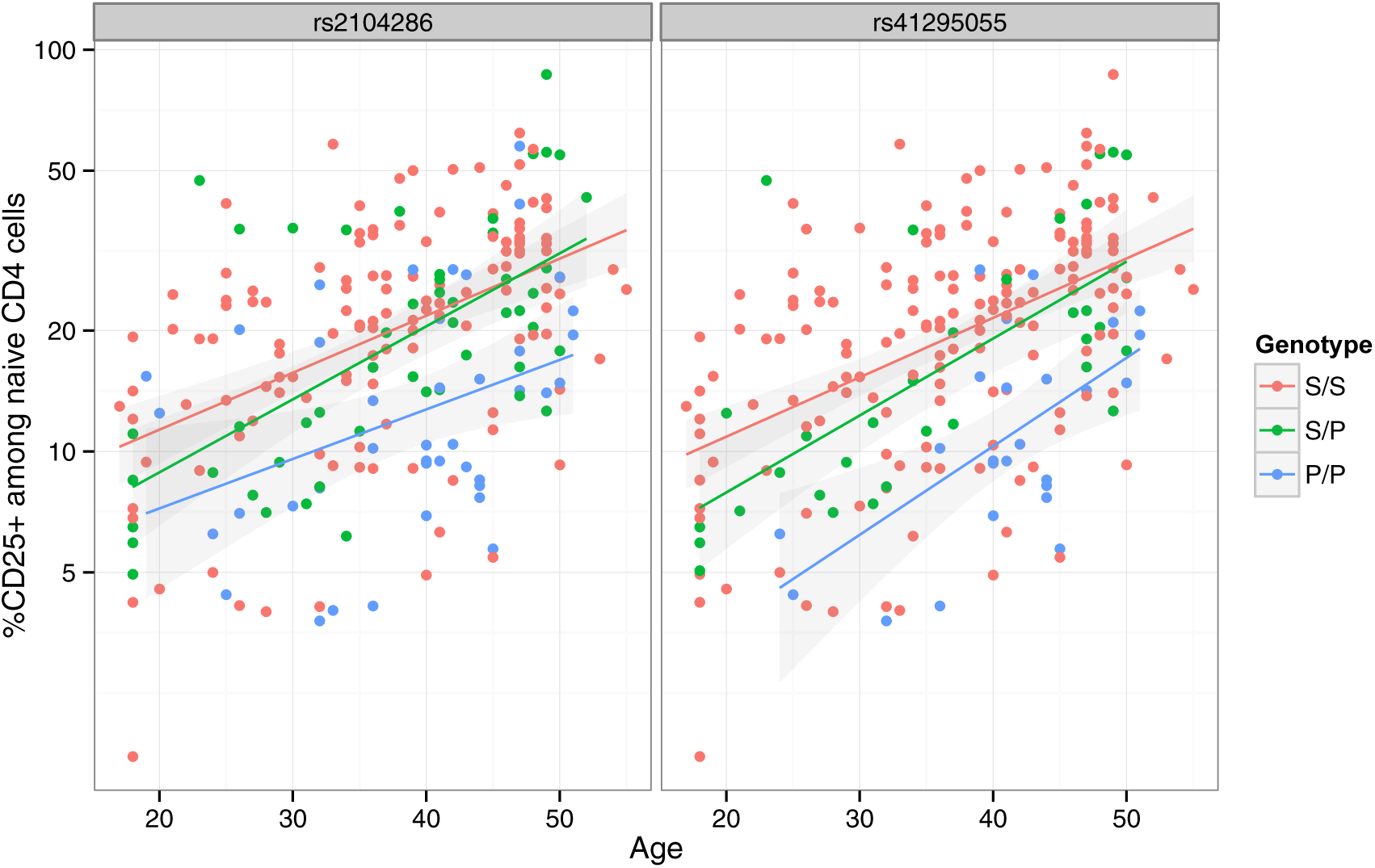
The proportion of naive CD4+ T cells that express CD25 (log scale) increases with age. The MS protective allele for the M2 SNP rs41295055:C>T associates with fewer CD4+ T cells expressing CD25 across all ages (*p* = 3.45 × 10^-8^), and is statistically preferred to the previously reported Ml SNP, rs2104286:T>C (*p* = 2.56 × 10^-6^; Δ BIC=8.43). S and P are used to represent the (common) MS-susceptible and (rare) MS-protective alleles respectively at each SNP. These SNPs are in limited LD (*r*^2^ = 0.3).

## Discussion

The 10p15 region contains at least five apparently distinct associations to the immune-mediated diseases, T1D and MS. Our results are the first to support a four SNP model in T1D, which most likely reflects a combination of increased sample sizes and the increased variant density available due to ImmunoChip and imputation, as this model was supported by both stepwise regression and GUESSFM. A previous comparative study of MS and T1D susceptibility in this region using conditional analysis of tag SNPs identified three groups of SNPs [26]: their group I matches our group A, their group II our group C, and their group III our group B. The results showed that the minor allele of SNPs in all these groups was protective for T1D, but that group I (A) was not associated with MS and that group II (C) was susceptible for MS. Our results, based on larger sample sizes and more extensive variant coverage, suggest that the minor allele of SNPs in group A, shared between T1D and MS, are associated with the same protective effect for both diseases. This emphasises the need for surveying the most complete set of variants possible in order to make cross disease comparisons.

Variants in this region have previously been linked to several aspects of IL-2R signalling in T1D and MS patients [27] and the MS-associated variants we identify (whether under the M1 or M2 models) all showed some evidence of association with expression of CD25 on the surface of T cells, linking *IL2RA* in a primary way into the etiology of this disease. For T1D, we were only able to link the *IL2RA* intron 1 signal, shared with MS under the M1 model, to CD25 expression on memory T cells. Neither the intergenic T1D SNP group (C), despite physical overlap with the MS associated SNP group D, indexed by rs41295055, nor the sets near *RBM17* and *PFKFB3* (E and F) have yet been linked to *IL2RA* expression. These signals could relate to CD25 expression on other cell subsets or under specific conditions. The cell-specific regulation of *IL2RA* expression and its role in modulating the immune system are both complex and only partially understood. For example, in addition to T cells, *IL2RA* is also expressed on many other cell types, especially under inflammatory conditions [28]. We have data on only a subset of IL-2R phenotypes that have been previously reported, and previous studies have genotyped only a subset of the disease associated SNPs described in this paper. Further genotyping of large samples with IL-2R related phenotypes is warranted to properly assess any other potential effects of the disease signals we identify on IL-2R signalling. Alternatively, it may be that other genes in the region, which have not been as well studied as *IL2RA,* are also causal for T1D, either directly or through interaction with *IL2RA.* For example, *PFKFB3*, an inducible 6-phosphofructo-2-kinase/fructose-2, 6-bisphosphatase isoform, allows rapid responses to energy requirements and insufficiency can lead to apoptosis and an inability to undergo autophagy in CD4+ T cells in RA [29]. *PFKFB3* has also been implicated in regulating insulin secretion in pancreatic beta cells [30], which could explain the T1D specific signals across this gene. All three genes, *IL2RA, PFKFB3* and *RBM17* are upregulated in CD4+ T cells upon *ex vivo* activation (Table S3) and two candidate SNPs, from groups E and F, sit between two DNase I sites close to the *RBM17* promoter. Furthermore, gene regulation can extend over 100 kb [31] and further studies will be needed to confirm the gene(s) through which these other SNPs influence T1D risk.

Two or more independent signals are commonly observed in fine mapped autoimmune disease regions using stepwise regression [7-10]. Despite long established doubts regarding the validity of stepwise regression [3], its use has continued to dominate in GWAS because of the number of SNPs measured simultaneously. Stepwise regression addresses the questions: ‘which is the single variant that can best explain the maximum trait variance?’ and, conditional on this single variant, ‘do any other variants explain additional trait variance?’ This is not equivalent to asking what set of variants best jointly explain trait variance. Often, as we have observed for T1D, stepwise regression may be expected to identify the same signals as a stochastic search approach. However, our analyses demonstrate that different results may be produced by stepwise and full multi-variant search approaches in some cases. In particular, we highlight and prefer a two SNP model for MS association to the *IL2RA* region, one of the SNPs shared with T1D (group A, indexed by rs12722495) and associated with CD25 expression on memory T cells, the other (group D, indexed by rs41295055) a better predictor for CD25 expression on the surface of naive CD4+ T cells than the SNP identified by stepwise regression (group C, rs2104286).

Methods that search the multimodal model space formed by multiple SNPs without conditioning on univarate association signals can therefore reveal a more accurate picture of the likely disease causal variants in any region. Other Bayesian [32], stochastic search [33] and penalised regression approaches such as lasso [34, 35] have been proposed on a GWAS scale, but these aim primarily to detect association in data sets with relatively sparse genotype coverage. Our focus is the adaptation of a Bayesian variable selection method to the fine mapping problem, and it is likely that an alternative method, such as piMASS [36], could have been substituted for GUESS in the model selection step. A detailed and direct comparison of piMASS with GUESS showed a similar recovery of models in simulated data, but indicated that GUESS was more computationally efficient [13]. We chose to use GUESS for its computation efficiency, but also because its g-prior formulation is specifically designed to deal with highly correlated predictors, a beneficial feature for fine mapping.

Applying the regularised regression approaches to our simulated data showed that the ROC curve for lasso with decreasing penalty out-performed stepwise analysis, but optimising the regularization parameter by minimising the cross validation error led to very high false discovery rates, suggesting that in this context this criterion is highly anti-conservative. Previous studies have also indicated that while regularised regression has strengths in terms of prediction, it may have weaknesses in terms of model selection [37], particularly for correlated predictors [38]. Elastic net [39] and the group lasso [40] have been proposed as regularised approaches tailored to correlated predictors, but in our hands did not outperform the simple lasso plus construction of a set of plausible SNPs according to *r*^2^ > 0.8. We considered that the information supplied to GUESSFM as a prior expected number of causal variants could be included equivalently in a regularised approach by setting the regularisation parameter according to the number of predictors selected. Although this produced a more competitive false discovery rate, discovery rates remained lower than with GUESSFM (Table S2).

Our multidimensional analysis strategy, tailored to the high genotyping coverage (and, hence, high LD) required for fine mapping causal variants can provide a more complete picture of the likely causal variants in a region. Any fine mapping study is limited by the set of variants included for study. While we attempted to survey the fullest possible set of SNPs and small indels by using dense genotyping data, plus imputation to the 1000 Genomes Project data, we cannot be sure we included all variants that might affect gene function without sequencing of cases and controls [41]. Further, larger indels, VNTRs and microsatellites remain particularly difficult to genotype with accuracy yet may contribute to disease. Therefore, it is important to bear in mind that all claims that a particular variant is causal are conditional on the true causal variants being accurately genotyped and included in the study.

It is well known that functional biological interactions may exist between variants [42], and therefore the possibility of statistical interactions needs to be considered in fine mapping studies. However, the need to fit interaction terms depends on the evolutionary history of the region. In the *IL2RA* region, there has been very little historical recombination (Figure S1) and *D′* between markers is nearly always 1, also indicating a lack of recombination [43]. As a consequence, only three of the possible four haplotypes that may be formed from any pair of SNPs are observed, and statistical interaction parameters are inestimable. This also implies that there is a one to one transformation between a haplotype model and a model expressing the log odds of disease as a linear function of single SNP effects [44], as we have employed. Thus, for the *IL2RA* region, we may neglect statistical interactions without making any assumptions about the existence of biological interactions. For our method to be applied to regions in which *D′* < 1, it would need extension to include statistical interaction terms. One simple approach, if the number of SNPs is not to large, would be to generate additional variables representing interactions between SNPs with *D′* < 1, but extending GUESS to fit interaction terms is another direction for further research.

An alternative approach would be to attempt to study disease risk across haplotypes directly [45], although these need to be inferred probabilistically when recombination has occurred. Haplotype-based analysis has become less common as marker density has increased. One reason for this is that haplotypes are particularly useful when marker density is low, because a haplotype may tag a causal variant better than any single variant, and haplotype studies have thus fallen out of fashion as denser genotype data have become available. This is exemplified by a study that found multiple statistical interactions underlying gene expression [46] (statistical interactions may be thought of as haplotype models) using SNPs on a genomewide array. Subsequent analysis found that all interactions which replicated in an external dataset with whole genome sequencing, and thus with more complete coverage of potential causal variants, could be better explained by a single SNP [47]. This suggests, perhaps, that statistical interactions between SNPs will prove rare. However, if biological interactions between variants separated by LD do exist, and the interaction depends on the phase of the variants, i.e., the diplotype risk differs from the two locus genotype risk, haplotype methods will again be required to infer likely causal variants.

One way to understand the means through which the effects of a disease-associated variant are mediated is to perform functional assays to examine its effects on a gene’s function [48] or to identify potentially intermediate phenotypes with which it is also associated [19]. Improved identification of likely causal variants should lead to more powerful and reliable follow-up studies, an important factor when many of these experiments require fresh primary cells and laborious wet lab protocols. Similarly, comparison with summary results from eQTL studies would be facilitated by the application of multi modal search strategies to the eQTL data set to ensure the effects of genetic variation are mapped as accurately as possible [49]. Even with large samples, and careful fine-mapping analyses, the causal candidacy of SNPs in high LD cannot be resolved through statistical association alone. Functional studies designed to directly address the confounding induced by linkage disequilibrium, such as allele-specific expression using rare haplotypes that distinguish SNPs in the same tag group, may be helpful in refining further the likely causal variants.

Informative approaches to understanding disease etiology have recently been developed based on looking for enrichment of the cell-specific chromatin marks localising to likely causal variants, and these are being used to highlight disease relevant cells [23,50]. Here, again, more accurate identification of causal variants will lead to more powerful and precise comparisons, and our approach, which is associated to a more accurate estimation of posterior probabilities for the SNPs in the considered region, should be readily adaptable to the growing set of methods that aim to examine enrichment [5] or colocalisation [6] by model averaging over the posterior probabilities that a SNP is causal.

## Materials and Methods

### Definition of target region

For fine mapping, we require dense coverage of genetic polymorphisms in the region. We targeted the ImmunoChip fine mapping region centred on *IL2RA,* namely chr10:6030000-6220000 (hg19). This region is bounded by recombination hot spots (Figure S1).

### Genotype data

Genotype data for this region comes from the MS [7] and the T1D [23] ImmunoChip studies. Quality control measures were applied to SNPs and samples as previously described [7]. As MS samples were derived from multiple international cohorts and we found allele frequencies for SNPs in the region varied between cohorts, we manually inspected plots of the first five principal components formed from ancestry informative markers [7] and additionally excluded samples lying outside the main cluster in each cohort.

### Imputation

We used IMPUTE2 [51] to impute untyped SNPs using the 1000 Genomes Phase I reference panel. We included a 500 kb window either side of our target region to allow variants in the less densely genotyped neighbouring regions to contribute information on the untyped SNPs.

### Adapted “shotgun” GUESS analysis

#### Tagging

Although GUESS is designed to work with correlated variables, the extreme, and occasionally perfect, LD in very dense genotyping data can lead to unstable results. We found, through a process of trial and error, that tagging so that no two SNPs remained with *r*^2^ > 0.99 retained good convergence of the stochastic GUESS algorithm. This reduced the number of SNPs from 667 to 443 (T1D) and 453 (MS). For each SNP that was removed, we noted its *index* SNP, the remaining SNP which had the highest *r*^2^ with it. We call a set of SNPs sharing a single index SNP a *tagset*; the number and size of tagsets is shown in Figure S6.

#### GUESS analysis

We ran GUESS in parallel on each disease using the index SNPs defined above. For MS, which included internationally derived samples, we included country of origin as a categorical covariate. Monitoring plots for each GUESS run, generated by R2GUESS [52] (http://cran.r-project.org/web/packages/R2GUESS), are shown in Figure S7 and Figure S8. We saved the top 30,000 models visited for each disease and for each saved model, we generated an expanded set of models, by adding each model that could be formed by replacing any index SNP with any of the SNP(s) in its tagset (Fig. 1). This expanded the set of models under consideration, M, ten fold, to 514,476. For each trait, we derived a posterior weight for each model in M. To be precise, we generated approximate Bayes factors for each model, *m ϵ M*, for each trait, using the Bayesian Information Criterion (BIC) approximation:

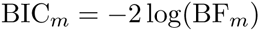

where

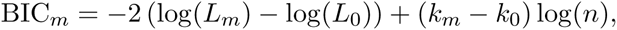

*L*_*m*_ and *k*_*m*_ represent the logistic likelihood of the data and the number of parameters under model *m, n* is the number of samples and 0 represents the null model. Finally we combined these Bayes factors with priors for each model, *π*_*m*_, to derive a posterior probability for each model

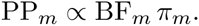

### Choice of priors

Priors could conceivably be generated on the basis of individual SNP content, for example, to prioritise those overlapping genomic annotations of particular interest. Such an approach has been adopted in a hierarchical framework, albeit with the single causal variant assumption [5]. For simplicity, and because we were interested to discover likely causal variants without predefining their likely mechanism, our model priors were determined only by the number of SNPs contained in a model, *N*_*m*_. A natural prior is then binomial or beta binomial. Given that, for a fixed expected value, a beta binomial puts larger weight on implausibly large models, we chose to use a binomial model, and, given published data on T1D, set the expected number of SNPs at three. For N SNPs in the target region, this means the prior for a model containing *N*_*m*_ SNPs is

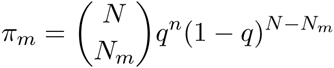

where *q* is set so the expected value of the binomial distribution was 3, ie 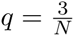

Priors and posteriors for the number of causal SNPs are shown in Figure S2.

#### SNP groups for candidate causal variants

We do not expect that we will be able to discriminate, statistically, between highly correlated variants. Instead, the posterior support is likely to be diluted across such sets of variants. To group such SNPs, we used the marginal posterior probabilities of inclusion (MPPI) for each SNP, and applied the following algorithm:

1. Pick the index SNP with maximum MPPI
2. Order remaining SNPs by decreasing *r*^2^ with index SNP
3. Exclude SNPs which co-occur in models with the index SNP (joint MPPI > 0.02)
4. Step away from the index SNP in order of decreasing *r*^2^, adding SNPs to its group until MPPI < 0.001 for two SNPs in a row (NB, these SNPs will not be added to the SNP group), or until *r*^2^ < 0.5
5. Remove this set of SNPs and return to step 1 until no SNP remains with MPPI > 0.01.

We summarize the support for any group of SNPs by the gMPPI, the sum of the posterior probabilities over all models containing a SNP in that group. The complete algorithm is available in the function snp.picker from the R package GUESSFM (https://github.com/chr1swallace/GUESSFM).

#### Model averaged effect estimates

We produced effect estimates for each SNP i as

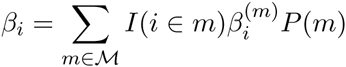

where 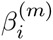 is the estimated effect of SNP *i* in model *m*, *I*(*i ϵ m*) is an indicator function taking the value 1 if SNP i is included in model m, and 0 otherwise, and *P*(*m*) is the posterior probability of *m ϵ M*.

#### Simulation analysis

We used simulated data to compare the performance of our proposed method, traditional stepwise regression, the lasso [53] and the elastic net [39]. For each simulation, we selected a random subset of 2,000 T1D control samples genotyped in the *IL2RA* region and a random set of between two and five “causal variants” from amongst the SNPs with MAF> 0.01. We simulated a Gaussian phenotype for which the causal variants acted in an additive manner to jointly explain 10% of the phenotypic variance. We conducted a total of 1,000 simulations for each scenario.

The data were analysed in parallel using the four different methods. We performed forward stepwise regression, and selected index SNPs at a given *p* value threshold, *α*, as those SNPs selected at any stage of the stepwise process with *p* < *α*. For each index SNP, we created pseudo “credible sets” as the set of SNPs with *r*^2^ > 0.8 with the selected index SNPs. Note, by this definition, simulated causal SNPs may appear in more than one SNP group. We calculated the false discovery rate as the proportion of SNP groups which did not contain a causal variant, and we calculated the discovery rate as the proportion of causal variants found in at least one selected SNP group.

We used the snp.picker function from GUESSFM (http://github.com/chr1swallace/GUESSFM) with default settings to define credible sets for causal SNPs. By definition of the algorithm, SNPs may only be members of at most one SNP group. We selected SNP groups according to varying thresholds for the gMPPI, and calculated false discovery and discovery rates as above.

For lasso and elastic net, we used the R package glmnet [54], and optimised the regularisation parameter *λ* by minimising the ten fold cross validation error using the `cv.glmnet()` function. For elastic net, we kept the folds constant across different values of *α ϵ* [0.1,1] and chose the pair of values (*α, λ*) which minimised the ten fold cross validation error overall. To examine the path of elastic net solutions, we fixed α at this value and varied λ. We used the R package gglasso [55] to implement the grouped lasso, predefining groups of variables as those with *r*^2^ > 0.8 with each other using heirarchical clustering. We also considered the discovery and false discovery rates with the first three or first five predictors selected, as something analagous to the prior information given to GUESSFM about our expectation of the number of causal variants.

#### Association with cell surface expression of CD25

Naive CD4+ T cells from 209 donors were gated as previously described [19, 56]. Association with index SNPs from disease-associated groups was assessed through linear regression. All possible one, two and three SNP models were considered, and the model with the minimum BIC reported. Expression was log transformed to reduce right skew of the phenotype.

#### DNase hypersensitivity data

We downloaded DNase I hypersensitivity for CD4+ T cells from the Roadmap Consortium [57] from http://vizhub.wustl.edu/VizHub/hg19, accessed 19 September.

#### RNA seq gene expression data

We measured gene expression in the studied region in two pooled samples (n=3 and n=4 individuals / pool) comparing unstimulated and stimulated CD4+ T cells. For each individual, 250,000 CD4+ T cells (93-99% pure, RosetteSep Human CD4+ T Cell Enrichment Cocktail, StemCell Technologies) were stimulated with anti-CD3/CD28 T-activator beads (Dynabeads Life Technologies) at a ratio of 0.3 beads / cell for four hours at 37°C in X-VIVO-15 (Lonza) + 1% AB serum (Lonza) and penicillin/ streptomycin (Life Technologies). Unstimulated CD4+ T cells were cultured in medium alone for four hours. Cells were harvested directly into Qiagen lysis buffer (Qiagen) and stored at -80°C until RNA isolation.

RNA was isolated using RNeasy micro kit including gDNA depletion (Qiagen). RNA integrity and concentration was evaluated using the Bioanalyzer platform (Agilent), with all samples showing an RNA integrity score (RIN) > 9.8. 750 ng of total RNA were used for the preparation of cDNA libraries using the Illumina TruSeq (Illumina) platform with a low-cycle-number PCR protocol, and was followed by transcriptome sequencing on an Illumina HiSeq 2000. This yielded four libraries with ∼38 million 100 bp paired-end reads each.

We trimmed raw reads to remove primer and adapter contamination, which affected 2% of our sequences, using HTSeq [58]. Reads were aligned to the reference genome Ensembl GRCh37.p13 using STAR [59]. Removal of low quality and unpaired reads, indexing, *IL2RA* region extraction, and depth counting were performed using SAMtools [60]. We employed HTSeq and DESeq2 [61] to carry out a differential expression analysis between the two conditions based on normalised read counts. We only considered paired-end reads that featured a total and unambiguous overlap with genomic sequence assigned to genes, around 73% of the initial raw sequences. Mapped read counts at each position in each sample are in Table S4.

This study was approved by local Institutional Review Boards (IRBs) and written informed consents were obtained from all participants in the study. For JDRF/Wellcome Trust Diabetes and Inflammation Laboratory the IRBs are: NRES Committee East of England - Cambridge Central (ref: 08/H0308/153), statistical analysis; and NRES Committee East of England - Norfolk (ref: 05/Q0106/20), for gene expression work. This study was approved by the University of Virginia Institutional Review Board (IRB number 17457); the Type 1 Diabetes Genetics Consortium collection sites obtained approvals for all subject collections and written consent and/or assent was obtained from all participants or their surrogates in the study.

## Acknowledgments

We thank all the T1D and MS patients and control subjects for participating in this study. We thank the members of each disease consortium who initiated and sustained the cross-disease ImmunoChip Project; Jeffrey Barrett for assistance with ImmunoChip SNP selection and for ImmunoChip related correspondence; Jennifer Stone for co-ordinating the ImmunoChip design and production at Illumina; Lorna Witty at the Wellcome Trust Centre for Human Genetics, Oxford for help in optimising cDNA library prep protocols and for performing the RNA sequencing. We thank the International Multiple Sclerosis Genetics Consortium (IMSGC) for sharing their ImmunoChip data. We thank David Dunger, Barry Widmer, and the British Society for Paediatric Endocrinology and Diabetes for the TID case collection.

We acknowledge use of DNA and RNA samples from the National Institute for Health Research (NIHR) Cambridge BioResource. We thank volunteers for their support and participation in the Cambridge BioResource and members of the Cambridge BioResource Scientific Advisory Board and Management Committee for their support of our study. Access to Cambridge BioResource volunteers and their data and samples is governed by the Cambridge BioResource SAB. Documents describing access arrangements and contact details are available at http://www.cambridgebioresource.org.uk/.

We acknowledge use of DNA from The UK Blood Services collection of Common Controls (UKBS-CC collection), which is funded by the Wellcome Trust grant 076113/C/04/Z and by the USA National Institute for Health Research program grant to the National Health Service Blood and Transplant (RP-PG-0310-1002). We acknowledge the use of DNA from the British 1958 Birth Cohort collection, which is funded by the UK Medical Research Council grant G0000934 and the Wellcome Trust grant 068545/Z/02. This research utilized resources provided by the Type 1 Diabetes Genetics Consortium, a collaborative clinical study sponsored by the National Institute of Diabetes and Digestive and Kidney Diseases, the National Institute of Allergy and Infectious Diseases, the National Human Genome Research Institute, the National Institute of Child Health and Human Development and the JDRF and is supported by the USA National Institutes of Health grant U01-DK062418.

The JDRF/Wellcome Trust Diabetes and Inflammation Laboratory is funded by the JDRF (9-2011-253), the Wellcome Trust (091157) and the National Institute for Health Research Cambridge Biomedical Centre. The research leading to these results has received funding from the European Union’s 7th Framework Programme (FP7/2007-2013) under grant agreement no.241447 (NAIMIT). The Cambridge Institute for Medical Research (CIMR) is in receipt of a Wellcome Trust Strategic Award (100140). CW is supported by the Wellcome Trust (089989). We acknowledge the National Institute for Health Research Cambridge Biomedical Research Centre for funding.

## Supporting Information

**Figure S1.**
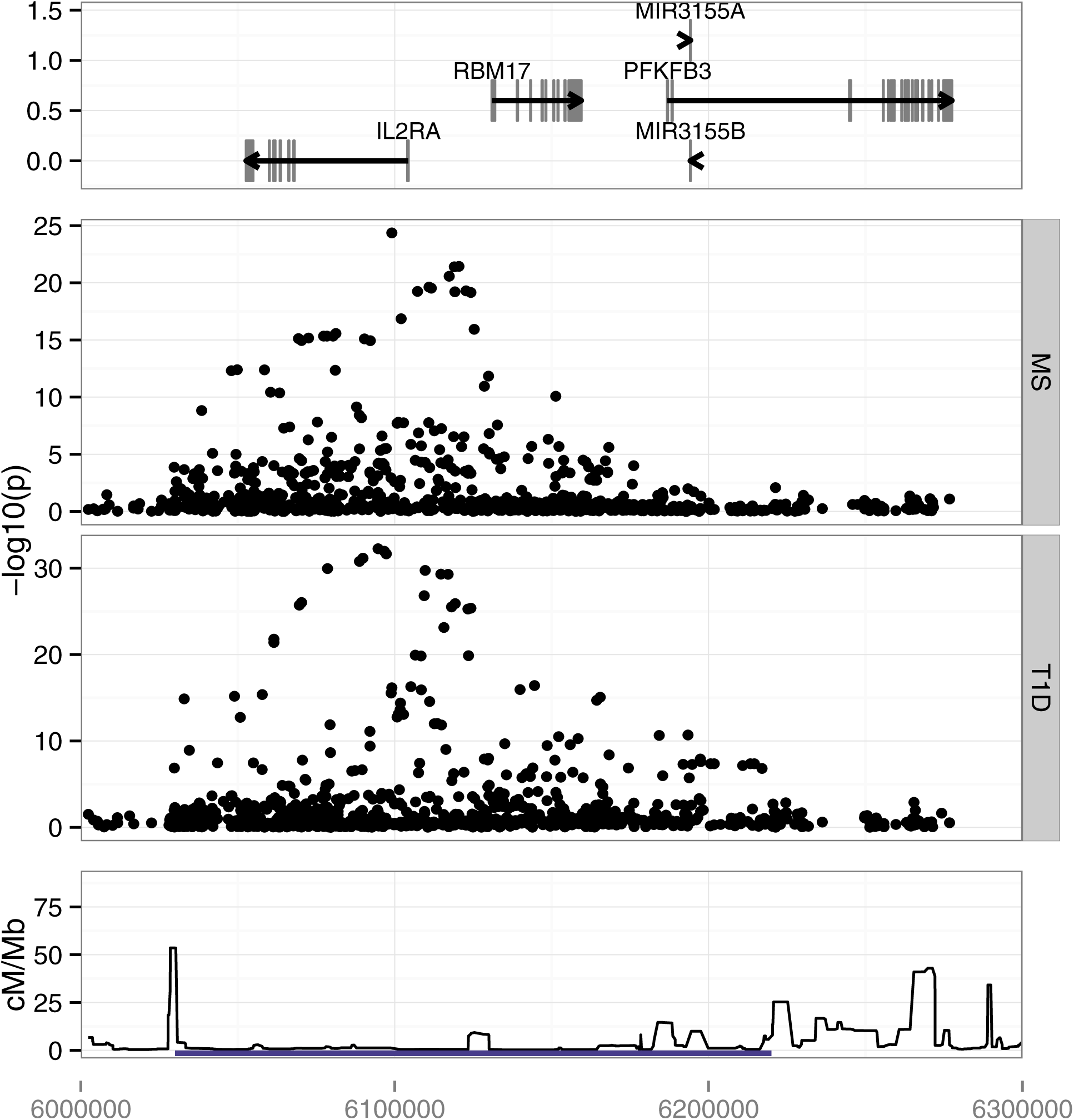
Manhattan plots for MS and T1D on chromosome 10:6,000,000-6,300,000 (hg19). Bottom track shows HapMap recombination rates and the blue bar indicates the region targeted for fine mapping.

**Figure S2.**
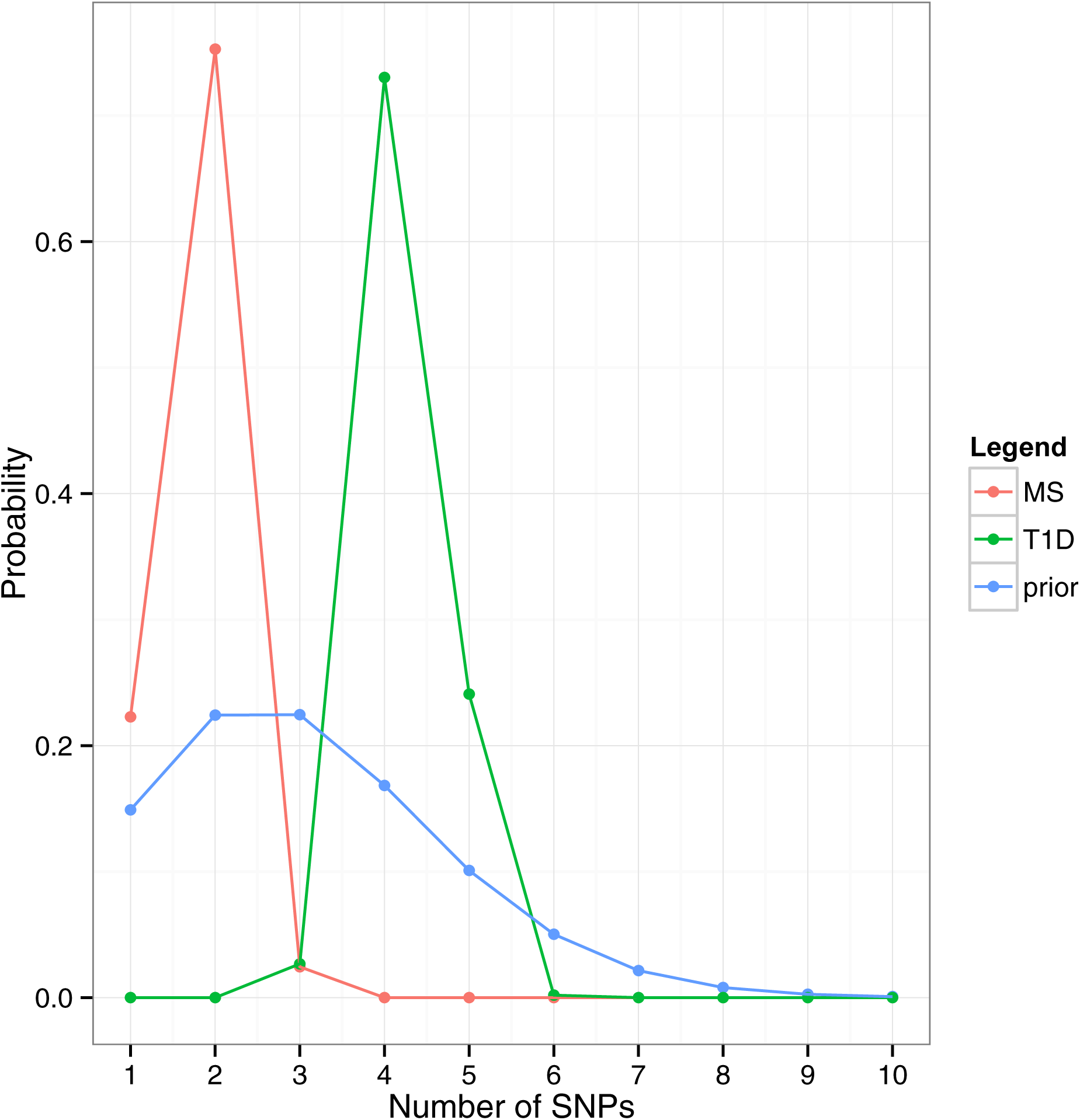
Prior and posterior probabilities across number of SNPs in underlying candidate causal models. For posteriors, this is the sum of posterior probabilities over all models visited by GUESS which contain the number of SNPs shown.

**Figure S3.**
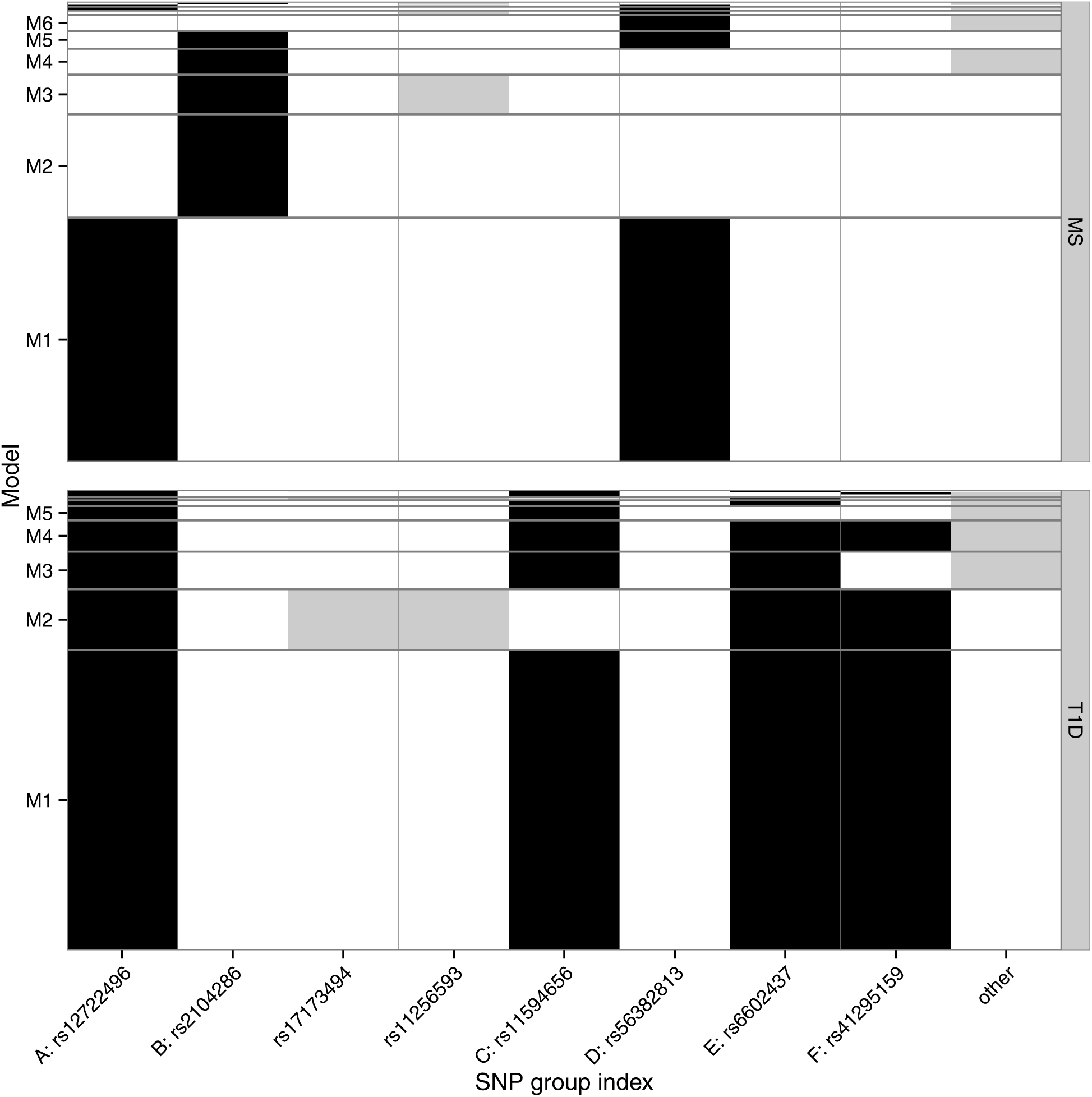
Patterns of inclusion of SNP groups in T1D and MS. Total filled bar height is proportional to gMPPI for the group indexed by the SNP shown on the × axis. Only the most probable SNP groups (gMPPI > 0.1) are labelled; “other” denotes SNPs outside these more probable SNP groups. Black fill indicates high confidence SNP groups (A-F) that were taken forward for further analysis. Groups indexed by rs17173494 and rs11256593 were considered to have too little support for either disease for association to be declared with confidence. For MS, we see two competing models: M1 indexed by A (rs12722496) and D (rs56382813), and M2, indexed by B (rs2104286).

**Figure S4.**
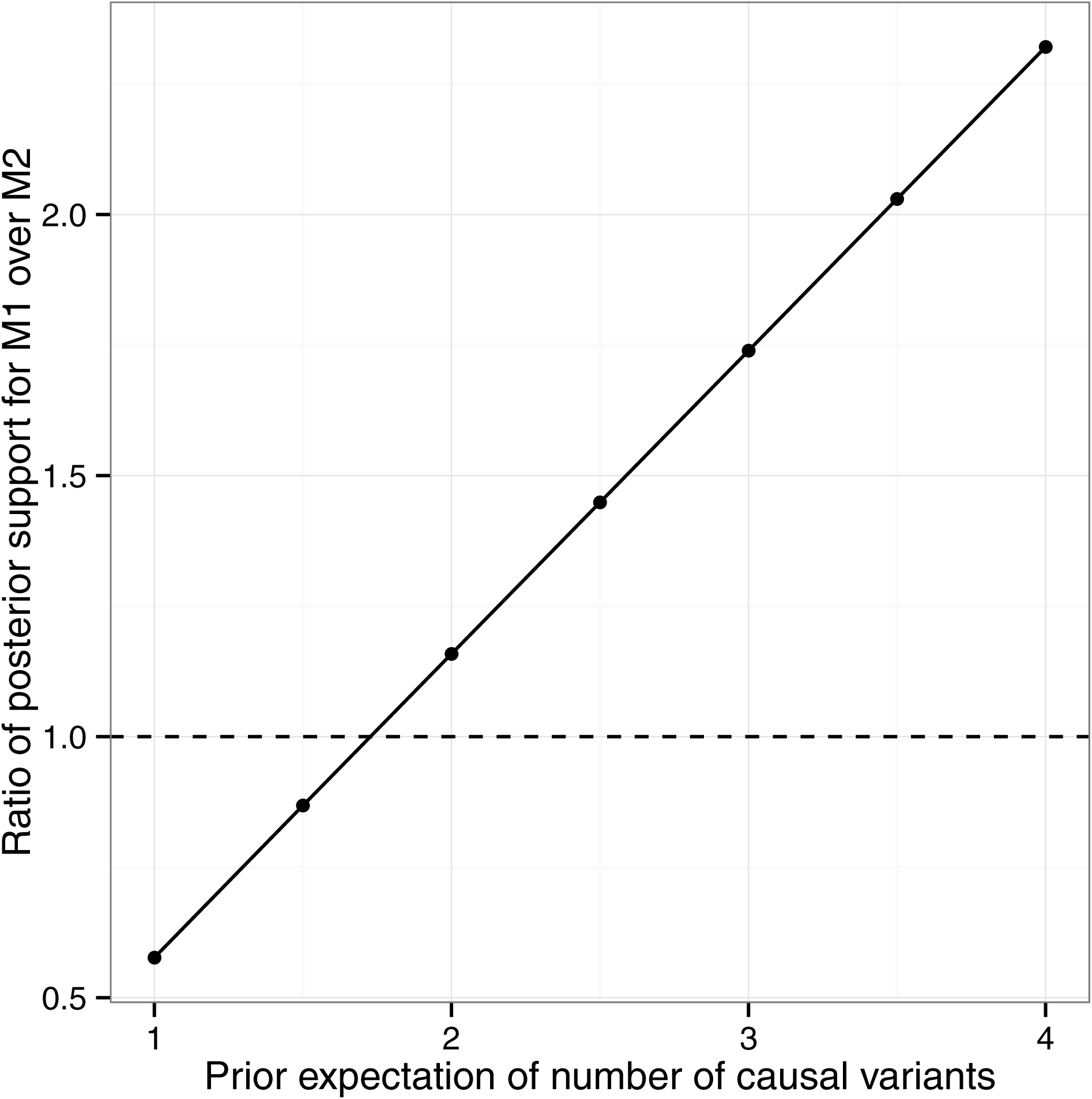
Sensitivity analysis showing the effect of varying the prior expectation of the number of causal variants on the relative posterior support for model M1 over M2 for MS. The relative posterior support is calculated as the posterior probability of all models within the M1 group divided by the posterior probability of all models within the M2 group for a given prior expectation. For a prior expectation of three, this is the ratio of bar heights for M1 over M2 from Figure S3.

**Figure S5.**
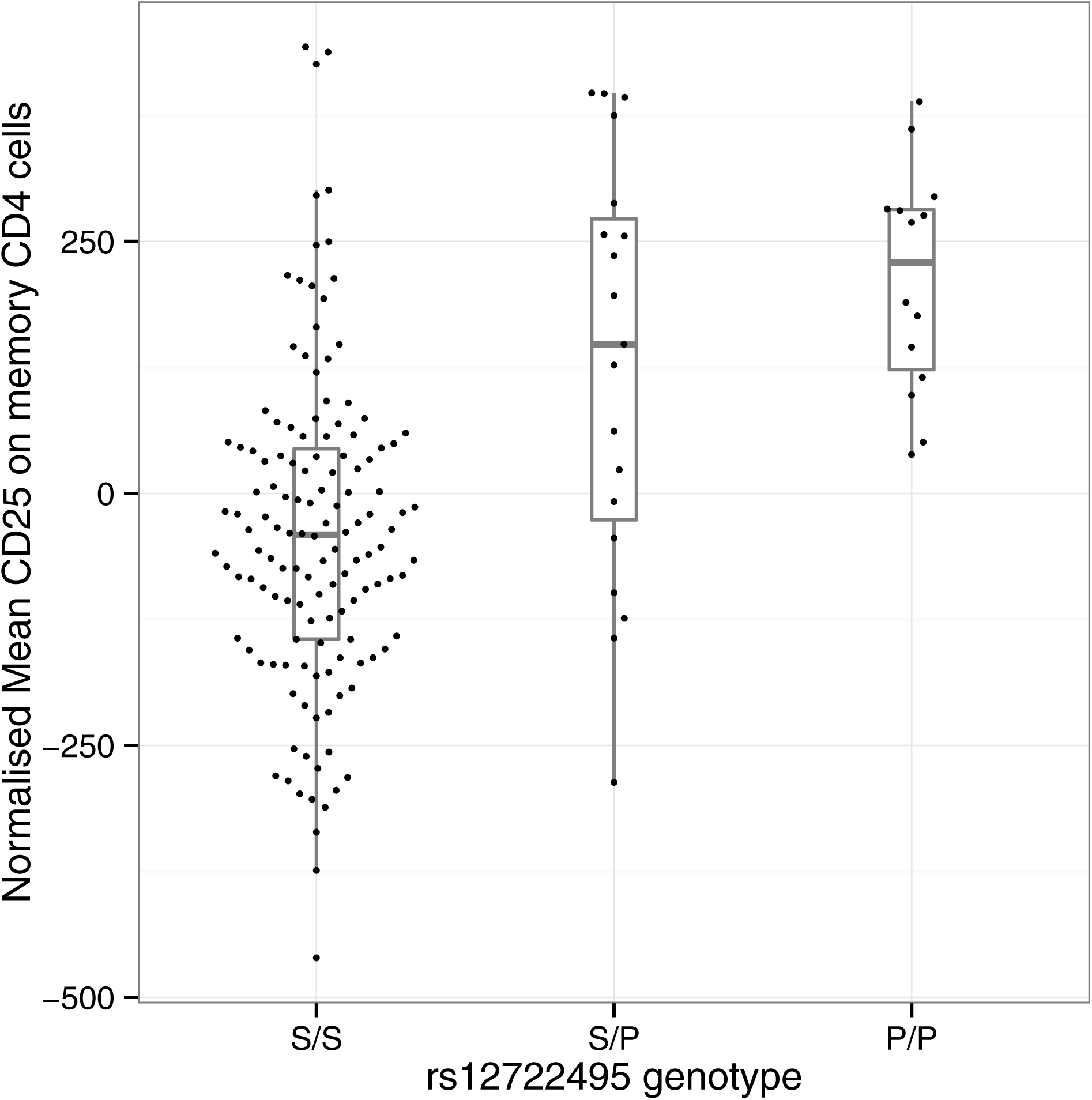
Normalised mean fluorescence intensity of CD25 dye on memory CD4+ T cells increases with copies of the (minor) T1D-protective (P) allele at rs12722495:T>C compared to the susceptible (S) allele, *p* = 5.50 × 10^-10^.

**Figure S6.**
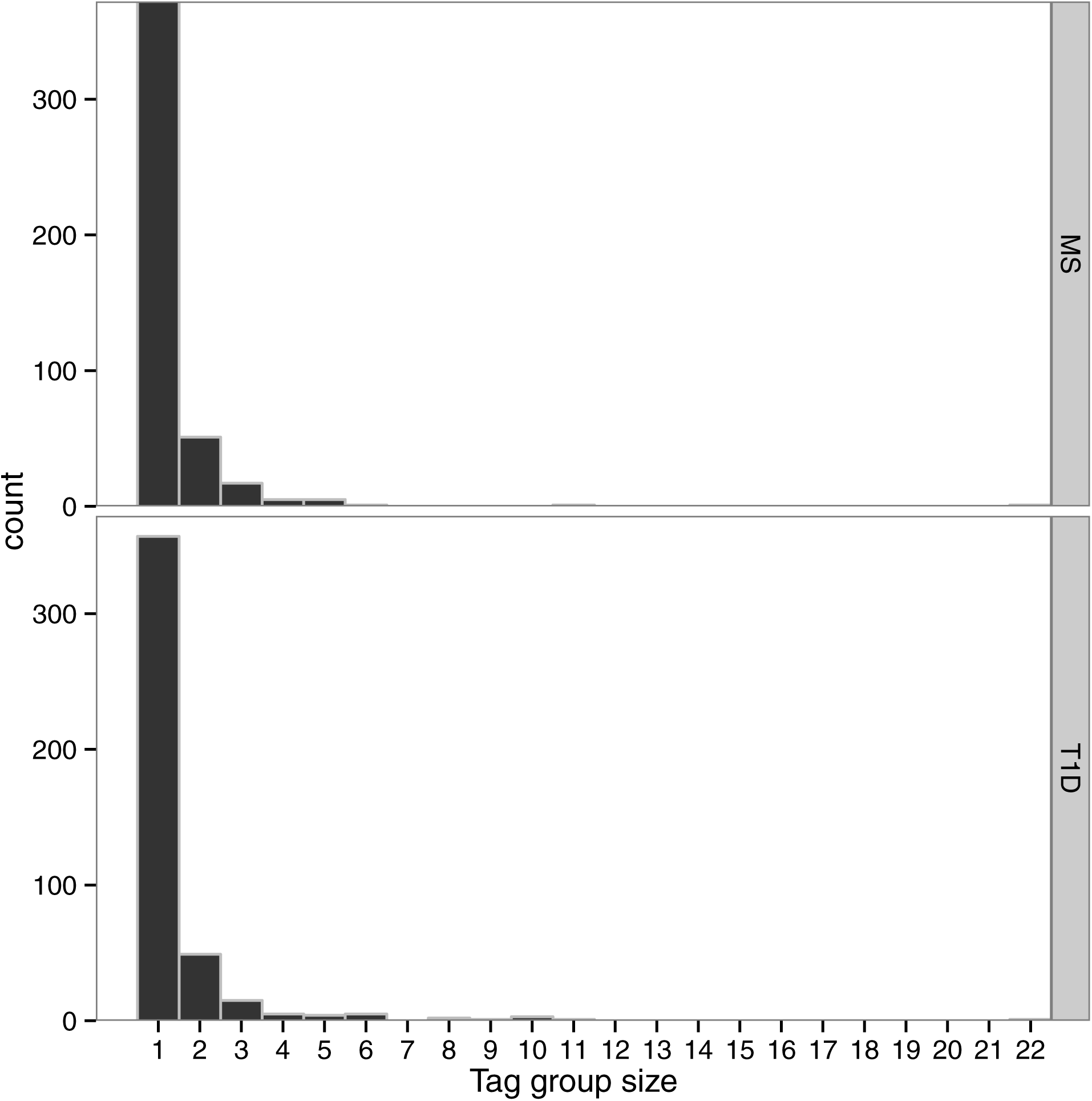
Size of SNP groups after tagging at *r*^2^ > 0.99.

**Figure S7.**
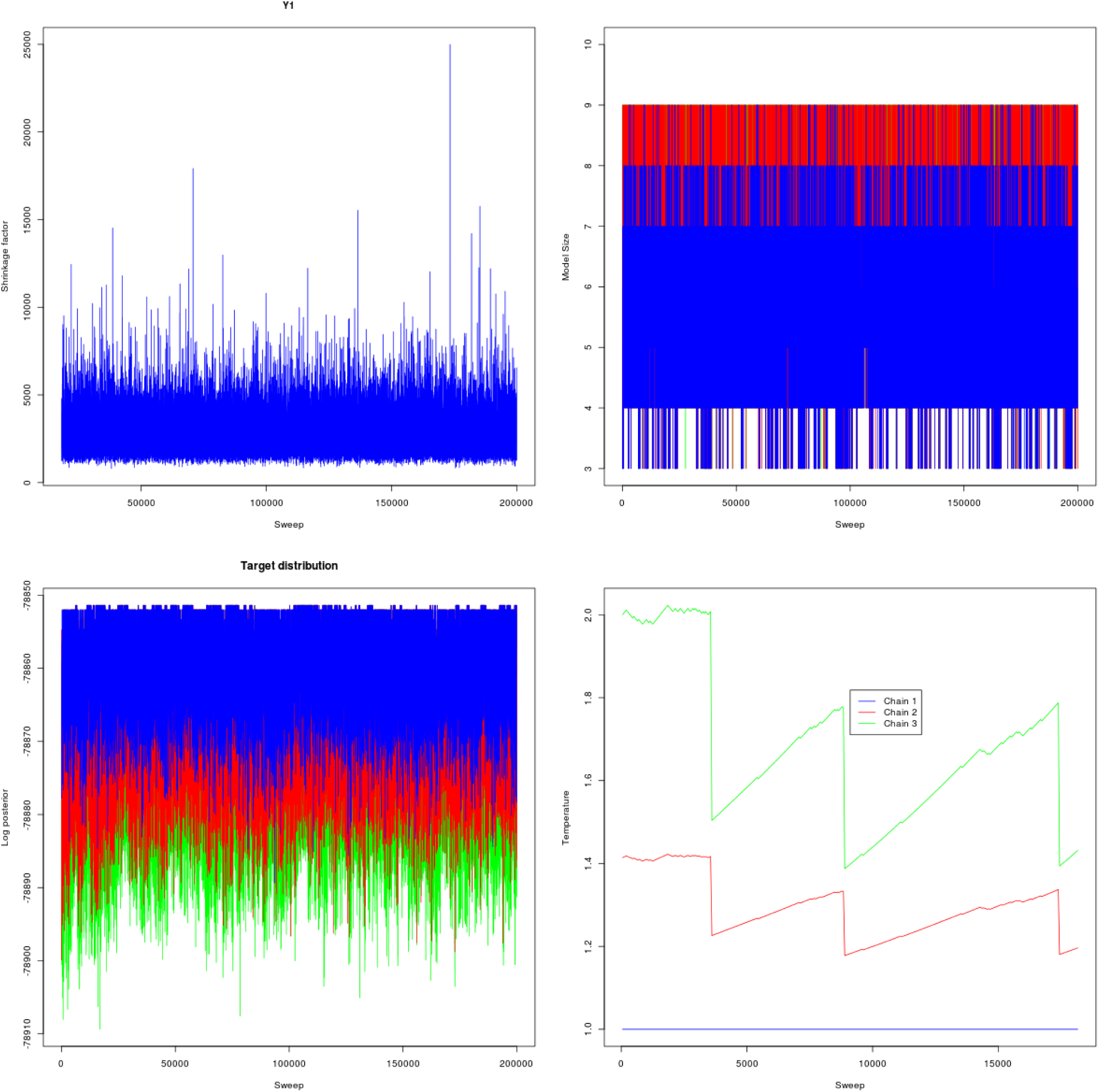
GUESS monitoring plots for T1D. The three chains are indicated by different colours.

**Figure S8.**
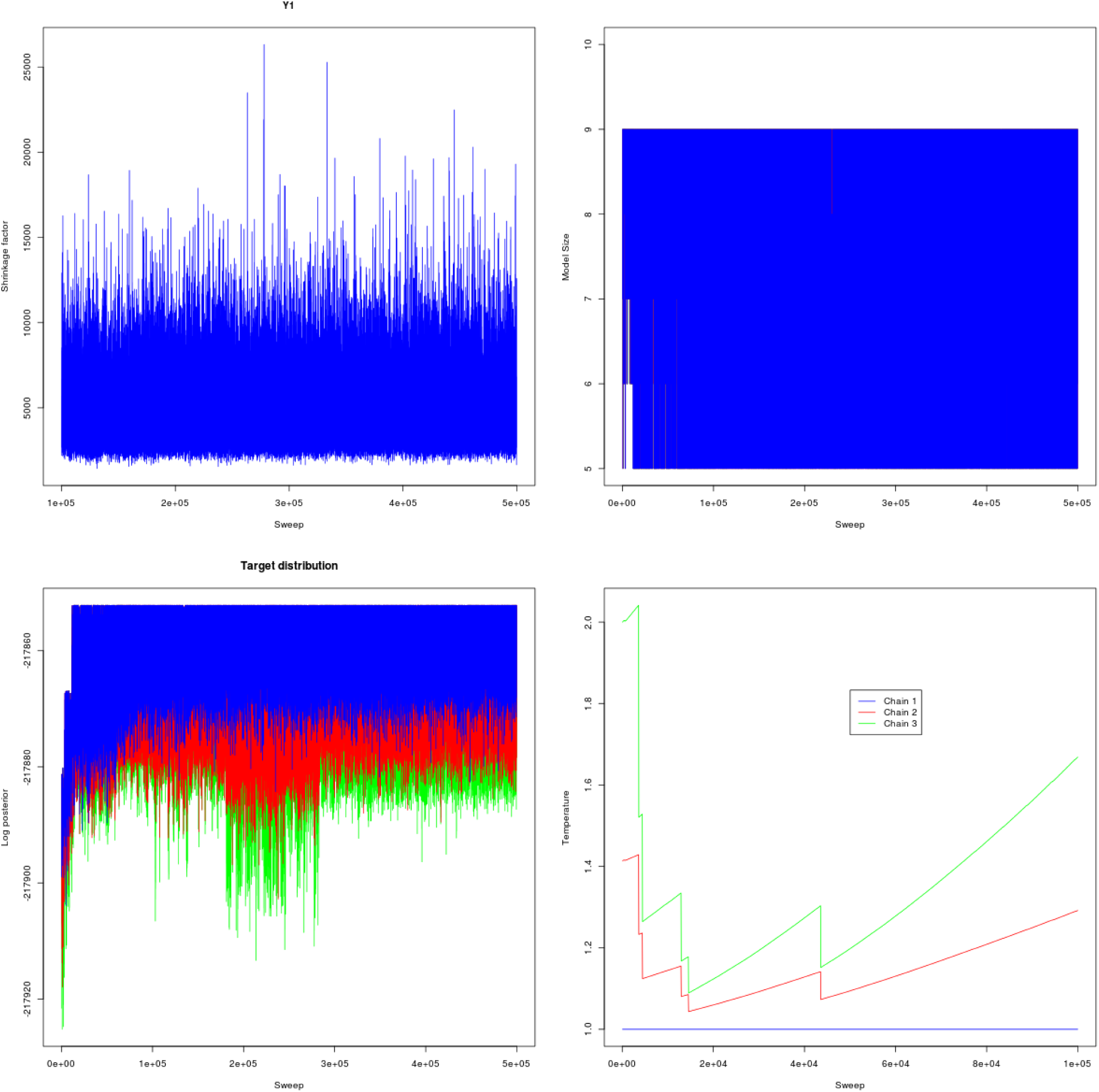
GUESS monitoring plots for MS. The three chains are indicated by different colours.

**Table S1.**
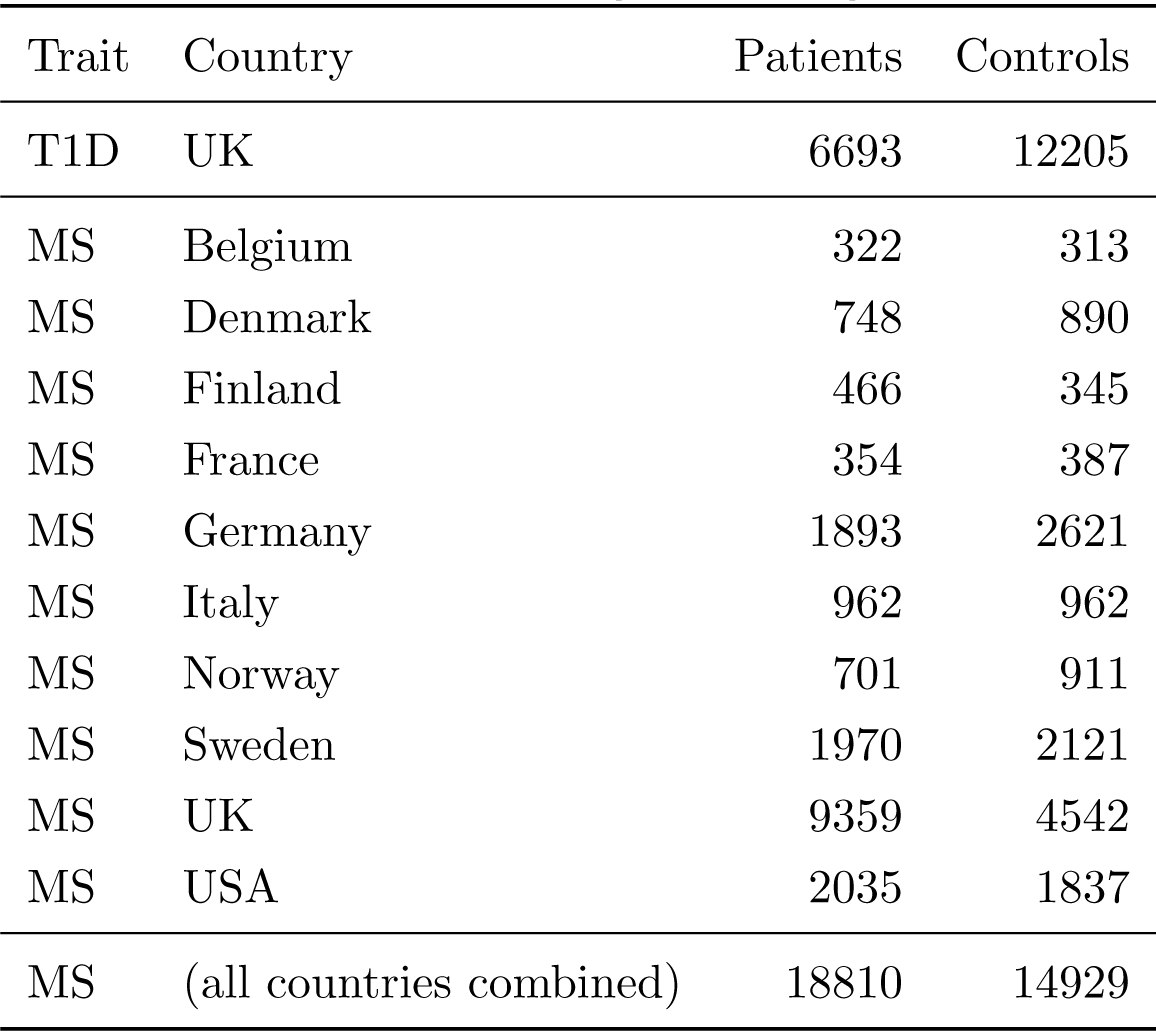
Samples with ImmunoChip genotyping data used in the fine mapping analysis.

| Trait | Country | Patients | Controls |
| --- | --- | --- | --- |
| T1D | UK | 6693 | 12205 |
| MS | Belgium | 322 | 313 |
| MS | Denmark | 748 | 890 |
| MS | Finland | 466 | 345 |
| MS | France | 354 | 387 |
| MS | Germany | 1893 | 2621 |
| MS | Italy | 962 | 962 |
| MS | Norway | 701 | 911 |
| MS | Sweden | 1970 | 2121 |
| MS | UK | 9359 | 4542 |
| MS | USA | 2035 | 1837 |
| MS | (all countries combined) | 18810 | 14929 |

**Table S2.** Discovery and false discovery rates in simulated data using different model selection methods and specified optimisation criteria, as described in Methods. Briefly, we ran GUESSFM setting the mean expected number of causal variants per region to either three (GUESSFM-3) or five (GUESSFM-5). We calculated the discovery rate (the proportion of causal variants within at least one credible set, y axis) and false discovery rate (proportion of detected variants whose credible sets did not contain any causal variant, × axis) at different thresholds for the stepwise *p* value (< 10^-6^ or < 10^-8^), the group marginal posterior probability of inclusion (gMPPI> 0.5 or > 0.9) for GUESSFM and the regularization parameter λ (chosen to minimise the ten-fold cross validation error, or at the largest value that selected exactly three or five predictors) across simulated datasets.

| Causal variants | Method | Criteria | Discovery rate | False discovery rate |
| --- | --- | --- | --- | --- |
| 2 | GUESSFM-5 | gMPPI $> 0.5$ | 0.966 | 0.009 |
| 2 | GUESSFM-5 | gMPPI $> 0.9$ | 0.939 | 0.003 |
| 2 | GUESSFM-3 | gMPPI $> 0.5$ | 0.970 | 0.015 |
| 2 | GUESSFM-3 | gMPPI $> 0.9$ | 0.943 | 0.004 |
| 2 | Lasso | min CV error | 0.994 | 0.982 |
| 2 | Lasso | first 3 predictors | 0.720 | 0.006 |
| 2 | Lasso | first 5 predictors | 0.880 | 0.015 |
| 2 | Stepwise | $p < 10^{-6}$ | 0.719 | 0.004 |
| 2 | Stepwise | $p < 10^{-8}$ | 0.644 | 0.003 |
| 2 | Elastic net | min CV error | 0.997 | 0.982 |
| 2 | Elastic net | first 3 predictors | 0.728 | 0.007 |
| 2 | Elastic net | first 5 predictors | 0.891 | 0.017 |
| 2 | Grouped lasso | min CV error | 0.986 | 0.947 |
| 2 | Grouped lasso | first 3 predictors | 0.671 | 0.021 |
| 2 | Grouped lasso | first 5 predictors | 0.846 | 0.044 |
| 3 | GUESSFM-5 | gMPPI $> 0.5$ | 0.900 | 0.011 |
| 3 | GUESSFM-5 | gMPPI $> 0.9$ | 0.797 | 0.002 |
| 3 | GUESSFM-3 | gMPPI $> 0.5$ | 0.900 | 0.021 |
| 3 | GUESSFM-3 | gMPPI $> 0.9$ | 0.833 | 0.005 |
| 3 | Lasso | min CV error | 0.992 | 0.974 |
| 3 | Lasso | first 3 predictors | 0.534 | 0.005 |
| 3 | Lasso | first 5 predictors | 0.692 | 0.013 |
| 3 | Stepwise | $p < 10^{-6}$ | 0.552 | 0.009 |
| 3 | Stepwise | $p < 10^{-8}$ | 0.458 | 0.005 |
| 3 | Elastic net | min CV error | 0.991 | 0.974 |
| 3 | Elastic net | first 3 predictors | 0.548 | 0.008 |
| 3 | Elastic net | first 5 predictors | 0.715 | 0.016 |
| 3 | Grouped lasso | min CV error | 0.964 | 0.937 |
| 3 | Grouped lasso | first 3 predictors | 0.481 | 0.019 |
| 3 | Grouped lasso | first 5 predictors | 0.648 | 0.040 |
| 4 | GUESSFM-5 | gMPPI> 0.5 | 0.856 | 0.023 |
| 4 | GUESSFM-5 | gMPPI> 0.9 | 0.747 | 0.008 |
| 4 | GUESSFM-3 | gMPPI> 0.5 | 0.801 | 0.033 |
| 4 | GUESSFM-3 | gMPPI> 0.9 | 0.690 | 0.014 |
| 4 | Lasso | min CV error | 0.995 | 0.967 |
| 4 | Lasso | first 3 predictors | 0.424 | 0.005 |
| 4 | Lasso | first 5 predictors | 0.567 | 0.011 |
| 4 | Stepwise | $p < 10^{-6}$ | 0.440 | 0.011 |
| 4 | Stepwise | $p < 10^{-8}$ | 0.340 | 0.007 |
| 4 | Elastic net | min CV error | 0.995 | 0.966 |
| 4 | Elastic net | first 3 predictors | 0.438 | 0.008 |
| 4 | Elastic net | first 5 predictors | 0.588 | 0.016 |
| 4 | Grouped lasso | min CV error | 0.928 | 0.931 |
| 4 | Grouped lasso | first 3 predictors | 0.385 | 0.018 |
| 4 | Grouped lasso | first 5 predictors | 0.530 | 0.037 |
| 5 | GUESSFM-5 | gMPPI> 0.5 | 0.755 | 0.033 |
| 5 | GUESSFM-5 | gMPPI> 0.9 | 0.635 | 0.008 |
| 5 | GUESSFM-3 | gMPPI> 0.5 | 0.715 | 0.038 |
| 5 | GUESSFM-3 | gMPPI> 0.9 | 0.589 | 0.014 |
| 5 | Lasso | min CV error | 0.993 | 0.958 |
| 5 | Lasso | first 3 predictors | 0.360 | 0.005 |
| 5 | Lasso | first 5 predictors | 0.494 | 0.011 |
| 5 | Stepwise | $p < 10^{-6}$ | 0.346 | 0.014 |
| 5 | Stepwise | $p < 10^{-8}$ | 0.252 | 0.007 |
| 5 | Elastic net | min CV error | 0.993 | 0.958 |
| 5 | Elastic net | first 3 predictors | 0.389 | 0.010 |
| 5 | Elastic net | first 5 predictors | 0.518 | 0.016 |
| 5 | Grouped lasso | min CV error | 0.911 | 0.918 |
| 5 | Grouped lasso | first 3 predictors | 0.326 | 0.017 |
| 5 | Grouped lasso | first 5 predictors | 0.455 | 0.036 |

**Table S3.**
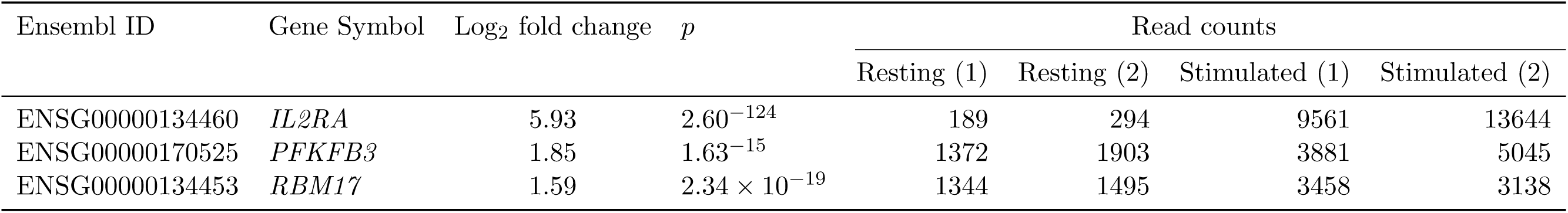
Differential expression analysis of genes in the region considered in this paper. All genes with measurable expression (reads > 20) in the studied samples are shown. Normalized read counts are shown for the two replicates of resting and stimulated CD4+ T cells.

Table S4

Mapped read counts in each sample in the target region from four RNA seq samples, as described in Methods.

separate file

